# Dirichlet distribution parameter estimation with applications in microbiome analyses

**DOI:** 10.1101/2024.04.17.589987

**Authors:** Daniel T. Fuller, Sumona Mondal, Shantanu Sur, Nabendu Pal

## Abstract

Microbiomes are of vital importance for understanding human and environmental health. However, quantifying microbial composition remains challenging and relies on statistical modeling of either the raw taxonomic counts or the relative abundances. Relative abundance measures are commonly preferred over the absolute counts to analyze and interpret microbiome (as the sampling fraction are unknown in sequence data) but currently there is no ideal distribution for carrying out this modeling . In this work, the Dirichlet distribution is proposed to model the relative abundances of taxa directly without the use of any further transformation. In a comprehensive simulation study, we compared biases and standard errors of two Methods of Moments Estimators (MMEs) and Maximum Likelihood Estimator (MLE) of the Dirichlet distribution. Comparison of each estimator is done over three different cases of differing sample size and dimension: (i) small dimension and small sample size; (ii) small dimension and large sample size; (iii) large dimension with both large and small sample size. We demonstrate the Dirichlet modeling methodology with four real world microbiome datasets and show how the results of the Dirichlet model differ from those obtained by a commonly used method, namely Bayesian Dirichlet-Multinomial estimation (BDME). We find that the results of parameter estimation can be dependent upon the sequencing depth and sequencing technique used to produce a given microbiome dataset. However, for all datasets, the Dirichlet MLE (DMLE) results are comparable to the BDME results while requiring less computational time in each case.

## 1 Introduction

### 1.1 Importance of microbiome study

Microbes are found in most environments on earth and microbiome depicts the diverse set of microorganisms that are present in an environment. Microbiome constitutes an important component of an ecosystem, maintaining its stability and sustainability through complex and dynamic interactions with plants, animals, and humans in the environment.[1, 2, 3] For example, soil microbes help in nutrient recycling and several species associated with plant roots are found to improve plant growth and protect them from pathogens.[1, 4] In marine environment, highly diverse life-forms exists at different depths and geographical locations, where a broad range of microbes help maintain a stable and healthy marine ecosystem through their vital contributions in various biogeochemical reactions and metabolic processes.[2] Distinct microbiomes are identified to be associated with different anatomical sites of the human body and play important roles in human health and disease.[3] In particular, interests in gut microbiomes has been rapidly growing in recent years with the complex communities of microbes inhabiting the gastrointestinal tract are found to be critical for intestinal homeostasis.[5] Gut microbiota establishes active bilateral interactions with host intestinal tissue and strongly influences host metabolism and immune system. A deviation from normal gut microbiome structure is reported to be associated with a number of disease conditions including inflammatory bowel disease, obesity, cardiovascular disorders, and even neurological disorders such as Alzheimer’s disease and Parkinson’s disease.[5, 6] Despite the importance of microbiome is increasingly recognized, one constraint for studying microbiome arises from the current technological limitations in their measurement, which restricts accurate inference on the microbial composition in an environment. To address this shortcoming, research communities are in search for appropriate statistical tools to improve the estimation of microbial composition.

### 1.2 Statistical background

The study of bacteria via methods such as 16S rRNA amplicon sequencing is neither experimental nor fully observational in nature, meaning that an assumption is made whenever the raw counts are assumed to be true observations of the environment. The raw counts of each taxon are bounded above by the sequencing depth which is independent of the actual microbial load in the environment from which the samples are drawn.[7] Additionally, the sampling fraction is unknowable, meaning that the raw counts in any given sample may not be representative of those present in the population of interest. Information about the population is only preserved in terms of proportions of each taxa within a sample, meaning that the counts must be standardized which creates a dataset that is compositional in structure.[8] That is, each sample sums to one and values represnt only the proportion of each taxa represented there. The difficulty presented in analyzing this type of data, often of high dimension, has resulted in the lack of a clear best-in-class approach. There are many methods for differential abundance analysis that provide different and even contradictory inferences on identical datasets, as shown in literature reviews by Weiss et al. (2017)[9] and Nearing et al. (2021).[10] The probabilistic methods for microbiome analysis can be separated into two broad classes: univariate methods used to model read counts of a single taxon such as Poisson and negative binomial models, and multivariate methods used to model read counts of multiple taxa such as Dirichlet-multinomial and nonparametric models.[11, 12, 13, 14, 15, 7]

The class of mulivariate models can further be expanded into those that are based on the analysis of raw counts and those that are based on the analysis of compositions. Bayesian Dirichlet-Multinomial models, for instance, start with the raw counts and make the assumption that the counts are multinomially distributed. This assumption is unreasonable because both the sampling fraction and the impact of amplicon replications are hidden for the analyst. Compositional models are those that properly ameliorate these issues, often through further transformations of the data. Some methods (e.g. ALDEx2 by Fernandes et al.) address the issues of compositional data by implementing common models such as the Baysian Dirichlet-Multinomial but utilize transformations to move the compositional data from simplex space to Euclidean space.[16]. Similar transformations into Euclidean space are commonly applied to compositional data in both microbiome analysis methods and in other fields where compositional data naturally occurs such as epidemiology and geology.[16, 17, 18] Mandal et al. (2015)[19] utilize similar transformations in the ANCOM method, but, they also apply an ANOVA model that tests all pairwise ratios of taxonomic abundances. Huang and Peddada (2020)[20] further improve this methodology by implementing a correction to bias due to sampling fraction. In contrast with previous methods, we aim to work directly with and only with relative abundance data without the use of any further transformations.

### 1.3 Dirichlet distribution in the context of microbiome analysis

In this work, we approach compositional data from an alternative perspective by assuming that the relative abundances of taxa are distributed according to the Dirichlet distribution, the multivariate generalization of the Beta distribution, without making any assumptions regarding the raw counts. A Dirichlet distribution of dimension *d* can fully capture the information about *d* different taxa and is the natural choice for a distribution on the unit simplex, the space where compositional microbiome data resides. Since the total number of taxa in the population is unknown, we assume the dimension *d* of the Dirichlet distribution to be the number of unique taxa found in all samples after cleaning and processing of the data. Starting directly with the Dirichlet distribution avoids many of the additional transformations and the stringent assumption of multivariate normality made by many existing techniques such as ANCOM. For a more in-depth discussion of the Dirichlet distribution and its varied applications, see the 2011 book by Ng, Tian, and Tang.[21] The Dirichlet distribution has previously been applied to model economic compositional data in the form of a Dirichlet general linear model.[22] While the Dirichlet distribution has been applied as a prior distribution for microbiome counts that are assumed to be multinomially distributed, it has not been previously been used as a direct model for microbiome relative abundances.[11]

### 1.4 Outline of the current study

The goal of this work is to provide a foundation for the use of the Dirichlet distribution for application in microbiome analysis. Through simulation, we quantify the effectiveness of several different point estimators for the Dirichlet distribution including cases (data with high dimension *d* but low sample size *n*) that commonly have increased false discovery rate for statistical tests.[20] There has never been a comprehensive simulation study run for these estimators that looks at realistic high *d* cases. Additionally, we provide applications of this work by estimating the Dirichlet parameter values relating to important taxa of interest in multiple gut microbiome datasets and compare the estimates against those of the Bayesian Dirichlet-Multinomial model proposed by Holmes, Harris, and Quince (2012).[23] The Bayesian Dirichlet-Multinomial model is popular and yet suffers from the issue of requiring count data to function. In order to ascertain the appropriateness of the Dirichlet distribution as a model for our microbiome datasets, we attempt to utilize a GoF test from Li (2015) to check the applicability of the model.[24] Unfortunately, the test is computationally difficult to apply for high *d* data and there has been no proper investigation of size and power for the test. Hence the GoF test for the Dirichlet distribution remains an open problem.

The manuscript is organized in the following manner. Section 2 provides basic information on the Dirichlet distribution and estimation of the Dirichlet parameter vector *δ* as well as the results of a simulation study comparing the merits of these different estimators. We consider the following cases: (i) small dimension and small sample size; (ii) small dimension and large sample size; (iii) large dimension. In Section 3, we investigate the asymptotic behavior of the MLE which provides some direction about what to expect for large sample sizes. Section 4 provides an example of fitting the Dirichlet distribution to four different real-world microbiome datasets and performing goodness-of-fit testing to ensure that the Dirichlet distribution is applicable.

The contributions of this work to existing literature are:

1. A comprehensive simulation study was performed for point estimators of the Dirichlet distribution
2. Wider ranges of dimension *d* and Dirichlet parameter vector *δ* were explored
3. The asymptotic properties of the Dirichlet MLE were investigated
4. The efficacies of Dirichlet point estimation and estimation via the Bayesian Dirichlet Multinomial model were compared with real world microbiome data

## 2 Estimation of the Dirichlet parameter vector

Parameter estimation is the primary means by which one can relate a distribution to data. There are several methods of parameter estimation available for the Dirichlet distribution including maximum likelihood estimation, method of moments estimation, and Bayesian estimation. Previous work on Dirichlet distribution parameter estimation has focused on the MLE and an MME but was limited in scope, studying the behavior of the estimators for a very small selection of *d* and *n*. See, for example, Dishon & Weiss (1980).[25] In this study, we will investigate the performances of the MLE and MMEs in a much broader setting compared to previous work.

### 2.1 Dirichlet distribution and basic properties

The Dirichlet family of distributions is the (*d*−1) dimensional analog of the Beta distribution, defining a family of unit sum-constrained probabilities or proportions on the (*d*−1) dimensional simplex, *C*^*d*^, where

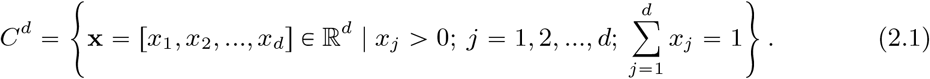

Suppose that a random vector X = (*X*_1_,*X*_2_, …,*X*_*d*_) ′, where 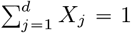 and *X*_*j*_ ∈ (0, 1), follows a (*d* −1) dimensional Dirichlet distribution with parameter vector ***δ*** = (*δ*_1_, *δ*_2_, …, *δ*_*d*_)′. Then the pdf of X is

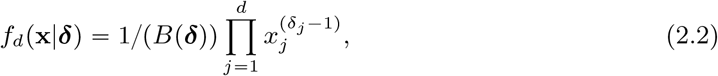

where *B*(·) is the Beta function, *δ*_*j*_ > 0 ∀ *j ∈* {1, 2, …, *d*}. The standard multinomial Beta function is

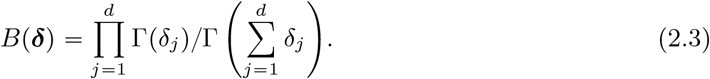

The Dirichlet distribution, henceforth referred to as Dir(***δ***|*d*), is not used directly in microbiome analysis as a distribution for relative abundances. It is commonly used, however, as a prior for multinomial models of microbiome count data. This set-up improperly assumes that the vector of raw (or absolute) counts of taxa follow a multinomial distribution, an assumption that our methodology skips (see the justification provided in Subsection 1.2). In Section 5 we estimate Dirichlet distribution parameter vectors from real-world microbiome data using the Bayesian Dirichlet-Multinomial method and compare it to estimation results using the MLE.

### 2.2 Point estimation

Suppose that 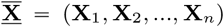 is a matrix of *n* independently and identically distributed *d*-dimensional microbiome observations from a population such that **X**_*i*_ ∼ Dir(***δ***|*d*).

#### 2.2.1 Method of Moments Estimator of the first kind (MME1)

The MME1 is used in various works in the literature and was originally used only as an MME for the Beta distribution.[26, 27, 21] We have adapted the MME from Ronning (1989), which is applied to the (*d* − 1) case, to the full *d* dimensional case. Let 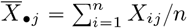 and <u>le</u>t 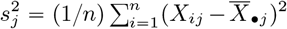, which is the sample variance of the jth component of 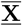. For *j* ∈ {1, 2, …, *d*},

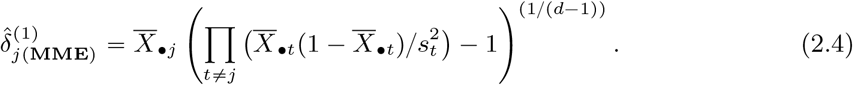

#### 2.2.1 Method of Moments Estimator of the second kind (MME2)

Let 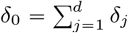 and let 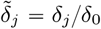. We find an alternate form of the MME by solving the following system of equations:

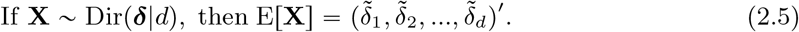

We can equate the above first population moment with the first sample moment, i.e.,

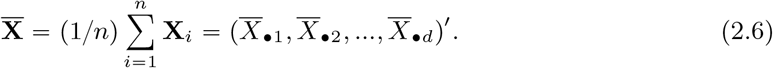

Note that due to the unit-sum constraint,

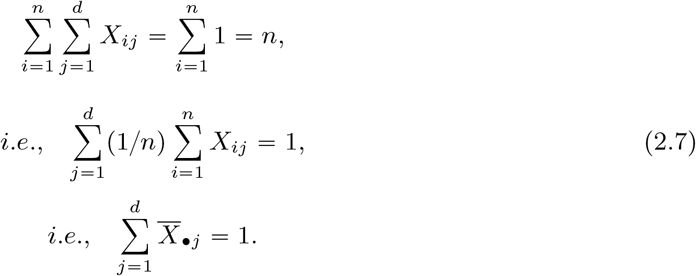

In order to obtain our estimator, our goal is to solve the following set of equations:

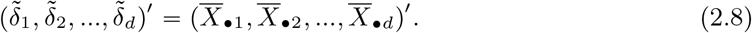

Suppose that X = (*X*_1_, *X*_2_, …, *X*_*d*_)′ ∼Dir(***δ***|*d*). Then,

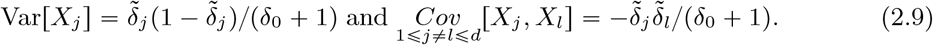

Set the sample variance 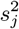 of the ith component of 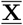 equal to Var[X_*j*_]. Equation 2.8 gives us that 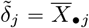. Then, from Equation 2.9, we get that

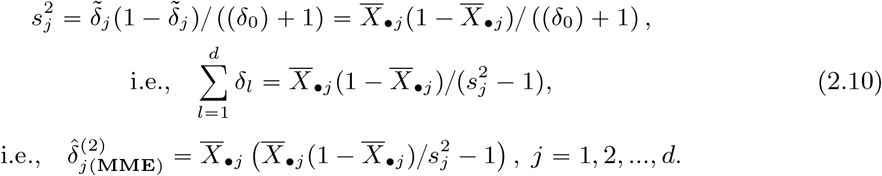

#### 2.2.3 Maximum Likelihood Estimator (MLE)

The most common estimator for the parameters of the Dirichlet distribution is the MLE, found by maximizing the log-likelihood, which is

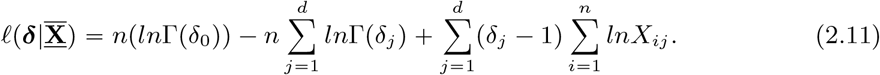

The MLE for *δ* is

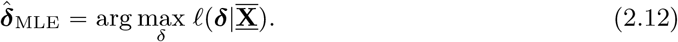

More information about the MLE for the Dirichlet distribution is available in Section 3. Previous work on Dirichlet parameter estimation has focused on the MLE and on MME1. Dishon & Weiss (1980) investigated the performance of 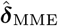 versus 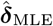 for the *d* = 2 and *d* = 3 cases of the Dirichlet distribution.[25] Since for large sample sizes the MLE should be the superior estimator,[28] the authors were interested in comparing the estimators over the sample sizes *n* = 25, 50, 100. Improvements to the initialization steps for algorithms for finding the MLE have been well-studied.[26, 29, 27] Despite that, to our knowledge no other work has directly compared the performance of different MMEs to the MLE. Additionally, this comparison has not been made for subspaces of the parameter space that accurately compare to the cases dealt with in microbiome data, namely high-dimensional data with a comparatively small sample size (*d* ¿¿ *n*).

### 2.3 Comparison of the estimators through a simulation study

In order to evaluate our estimators, we aim to generate 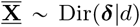 for a given parameter vector *δ*. First, generate *d* components Y_*j*_ such that Y_*j*_ ∼ Gamma(*δ*_*j*_, 1), *j* = 1, 2, …, *d* (with shape parameter *δ*_*j*_ and unit scale parameter). Define X_*j*_ as 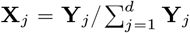 . Then 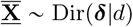.

For the *mth* set of replicated observations on X (1 ⩽ *m* ⩽ *M*), let the value of 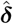 be equal to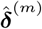. Define the estimation error vector as 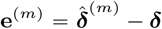. Then the bias, mean absolute error vector (MAEV), and mean squared error matrix (MSEM) are as follows:

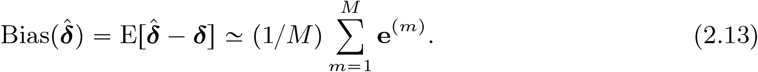

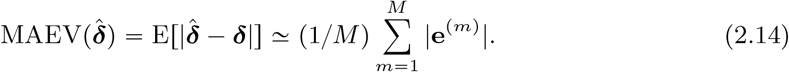

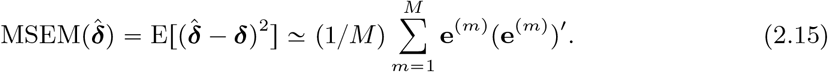

All simulations were run for *M* = 20, 000 replications. The dimension *d* was taken from *d* = 2 to *d* = 50. For brevity, results are shown for *d* = 2, *d* = 5, and *d* = 50. Sample size *n* was taken from *n* = 5 to *n* = 50 in increments of 5 and then from *n* = 50 to *n* = 300 in increments of 50. For the symmetric Dirichlet distribution, where *δ*_*j*_ = *δ* for all *j*, the common *δ* was taken in the range of 0.5 to 10. For the asymmetric case when *d* ⩽ 5, we tested choices of parameter values from *δ*_*j*_ = 0.5 to *d*_*j*_ = 10. For the asymmetric case when *d* > 5, *δ*_*j*_ are assumed to be independently and identically distributed Gamma random variables such that *δ*_*j*_ ∼ Gamma(*η*, 1). For paucity of space, we are presenting only representative tables and figures. The trends observed hold for other tables and figures not shown.

#### 2.3.1 Small dimension *d* and small sample size *n*

Bias and MAE for 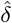 with small sample sizes and dimension *d* = 2 and *d* = 5are shown in Table 1. Bias and MAE over a range of *d* values are displayed in Figure 1 and Figure 2. Small sample Bias and MAE for estimated expected relative abundance are given in the appendix in Table 5.

**Table 1:**
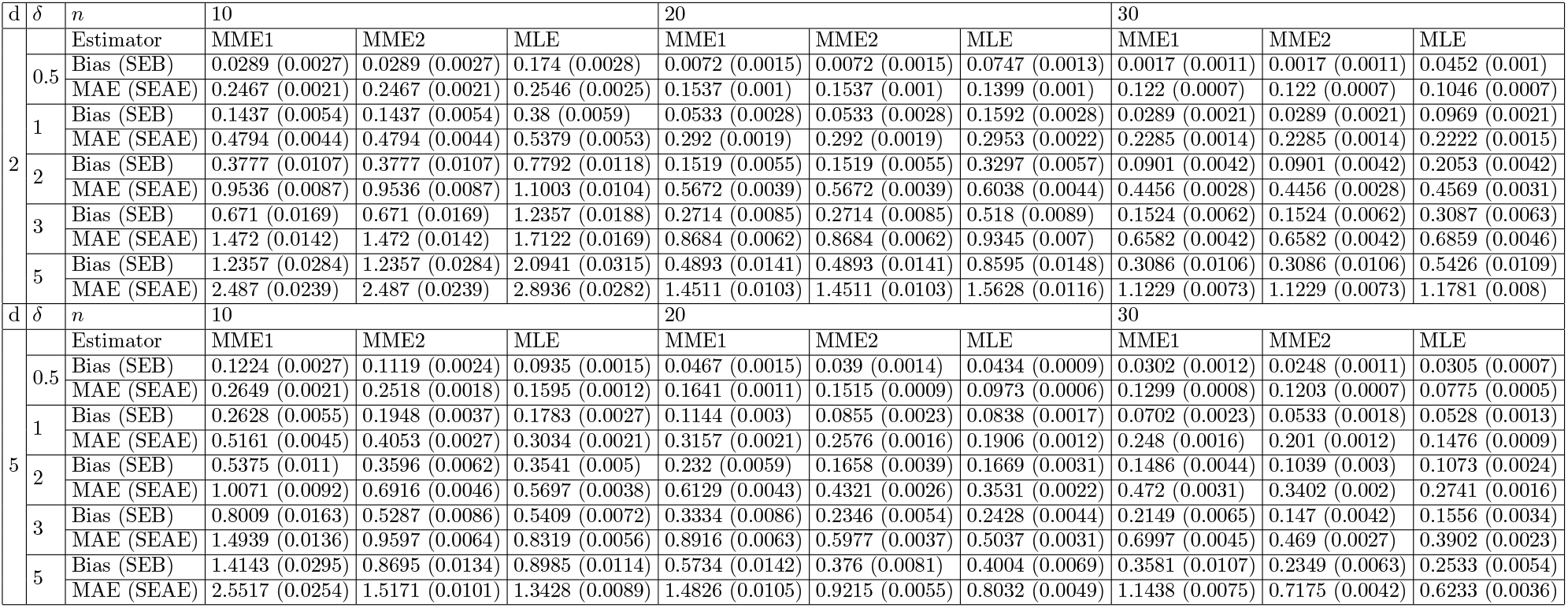
Small dimension, small sample estimation results for the symmetric Dirichlet with *δ ∈* {0.5, 1, 2, 3, 5}, *d* = 2, 5; *M* = 20, 000.

**Figure 1.**
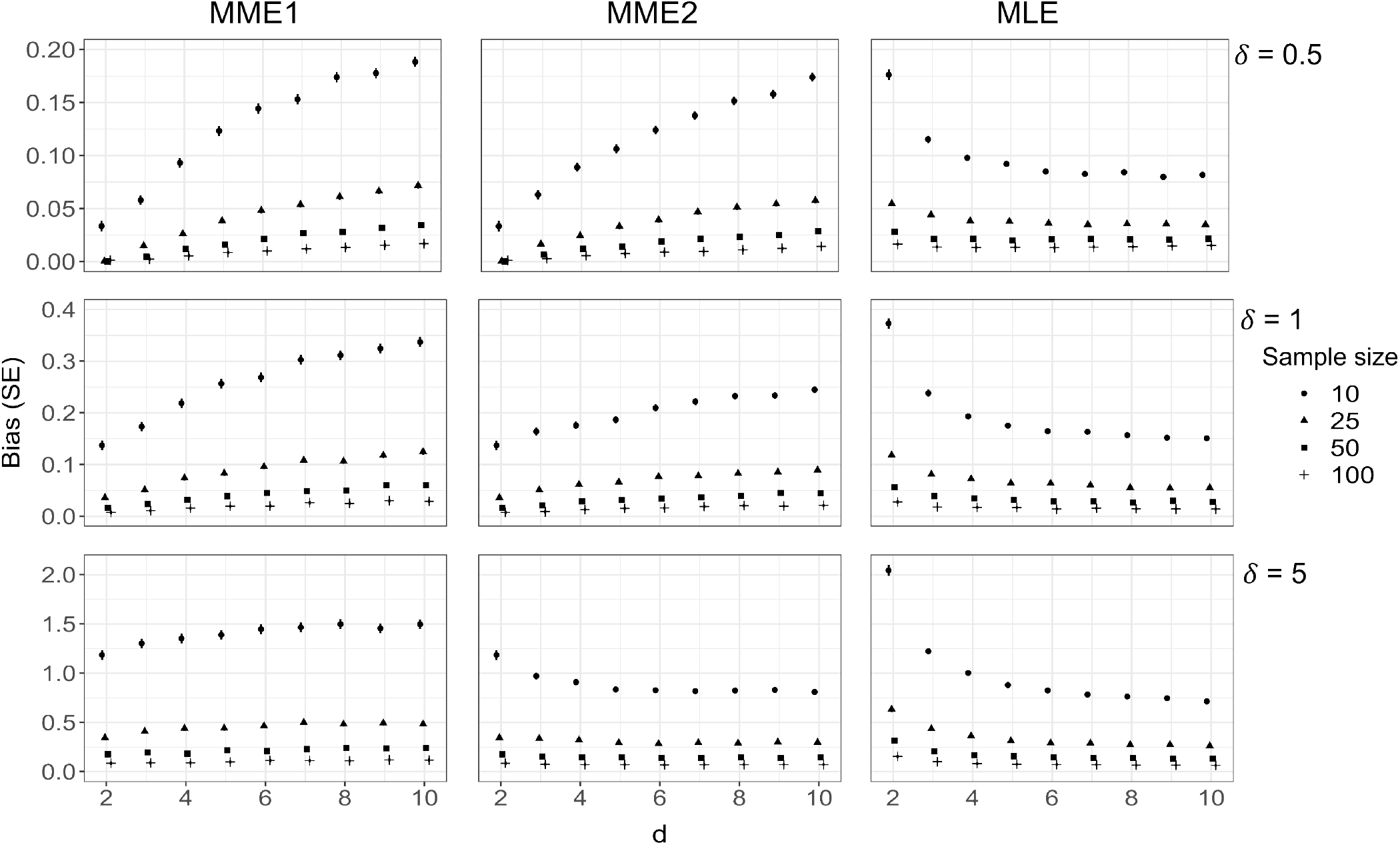
Estimation results for the symmetric Dirichlet distribution. Bias 1 ± SE are plotted against dimension *d* for all three estimators for sample sizes of 10, 25, 50, and 100. The common parameter value *δ* is taken to be 0.5, 1, and 5. The number of simulations, *M*, is 20,000.

**Figure 2.**
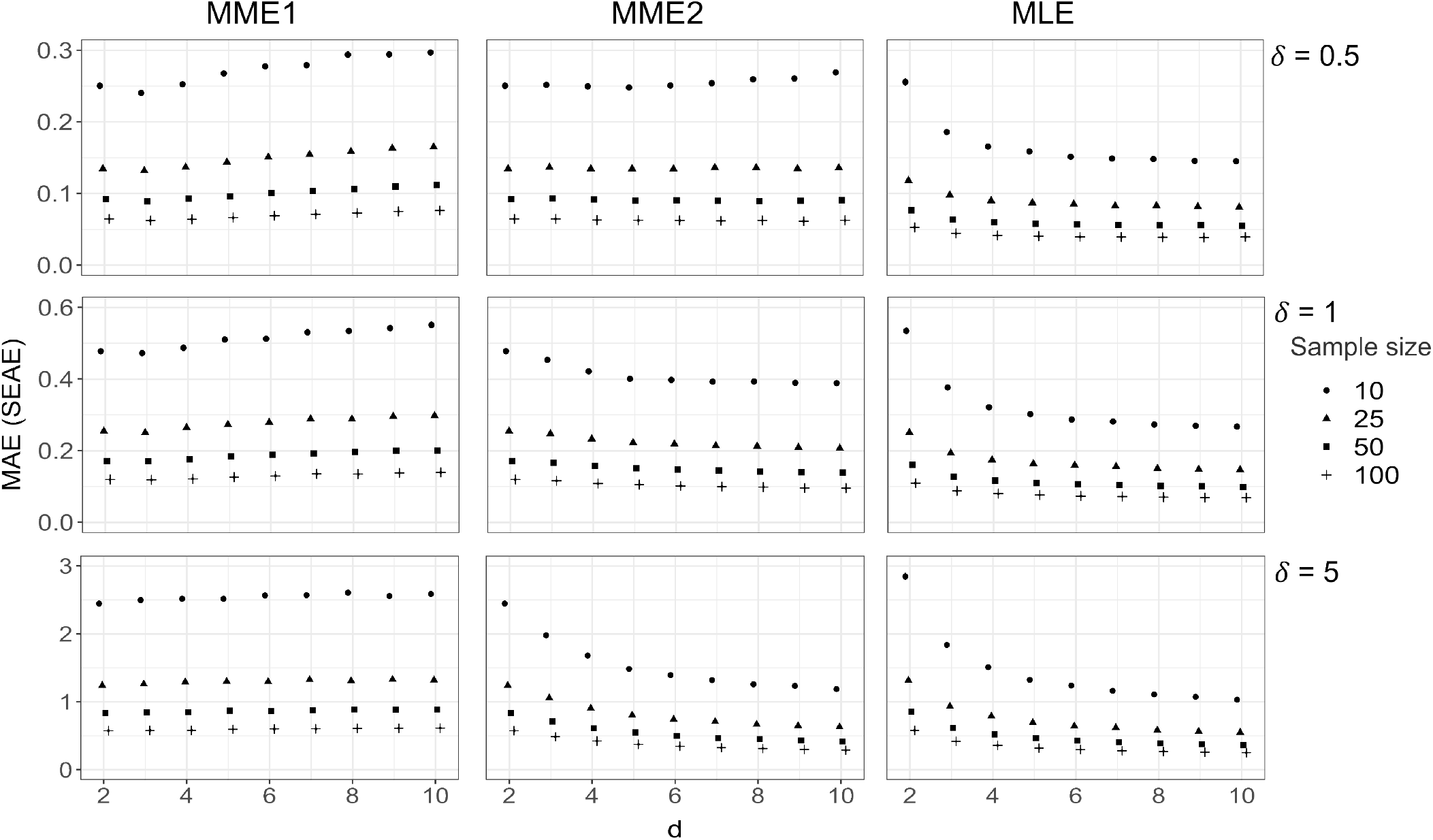
Estimation results for the symmetric Dirichlet distribution. MAE ± 1 SEAE are plotted against dimension *d* for all three estimators for sample sizes of 10, 25, 50, and 100. The common parameter value *δ* is taken to be 0.5, 1, and 5. The number of simulations, *M*, is 20,000.

#### 2.3.2 Small dimension *d* and large sample size *n*

Bias and MAE for 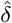 with large sample sizes and dimension *d* = 2 and *d* = 5 are shown in Table 2. Bias and MAE over a range of *d* values are displayed in Figure 1 and Figure 2. Large sample Bias and MAE for estimated expected relative abundance are given in the appendix in Table 6. As shown in Figure 1 and Table 2, the results for sample size *n* = 50 and higher are very similar compared to the results for lower sample sizes. This indicates that *n* = 50 appears to be the threshold between small and large sample sizes for these estimators.

**Table 2:**
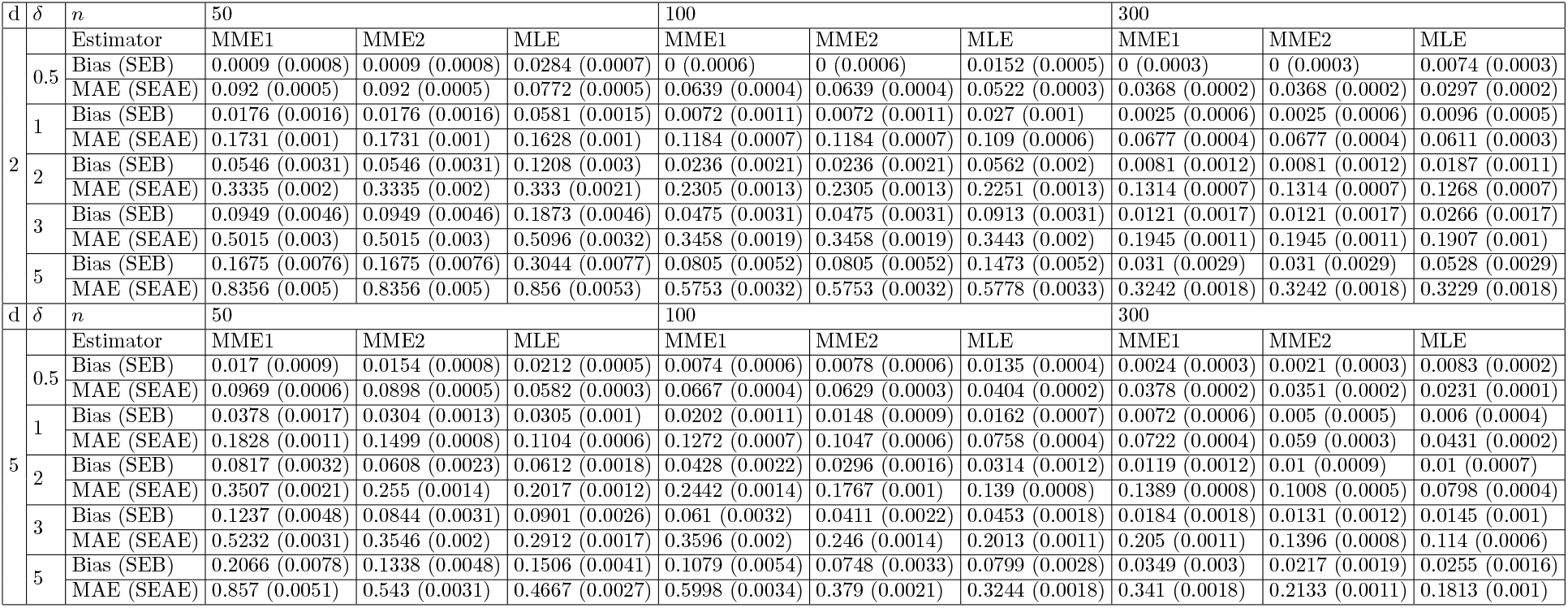
Small dimension, large sample estimation results for the symmetric Dirichlet with *δ* ∈ {0.5, 1, 2, 3, 5}, *d* = 2, 5; *M* = 20, 000.

Remark 2.1: As shown in Table 1, MME1 and MME2 exhibit very similar patterns of behavior for the symmetric case of the Dirichlet distribution. While their estimates diverge for smaller values of *n*, they are almost identical for large *n* as seen in Table 2. For both small and large sample sizes, the MMEs are less biased than the MLE although they tend to have larger standard errors, especially at small sample sizes. In terms of MAE, however, the MLE is either even with or outperforms the other estimators across sample sizes and parameter values. The difference in bias is much larger for *d* = 2 than *d* = 5. For higher dimensions, MME2 and the MLE both show comparatively better performance compared to MME1.

#### 2.3.3 Large dimension *d*

Tables 3 shows bias and MAE for both small and large sample sizes. Figure-3 shows bias for both small and large samples (*n* = 10, *n* = 25, and *n* = 50) when *d* = 50 for the asymmetric case of the Dirichlet distribution. Dirichlet parameter values are taken to be iid Gamma(*η*, 1). We present the results for *η* = 5.

**Table 3:**
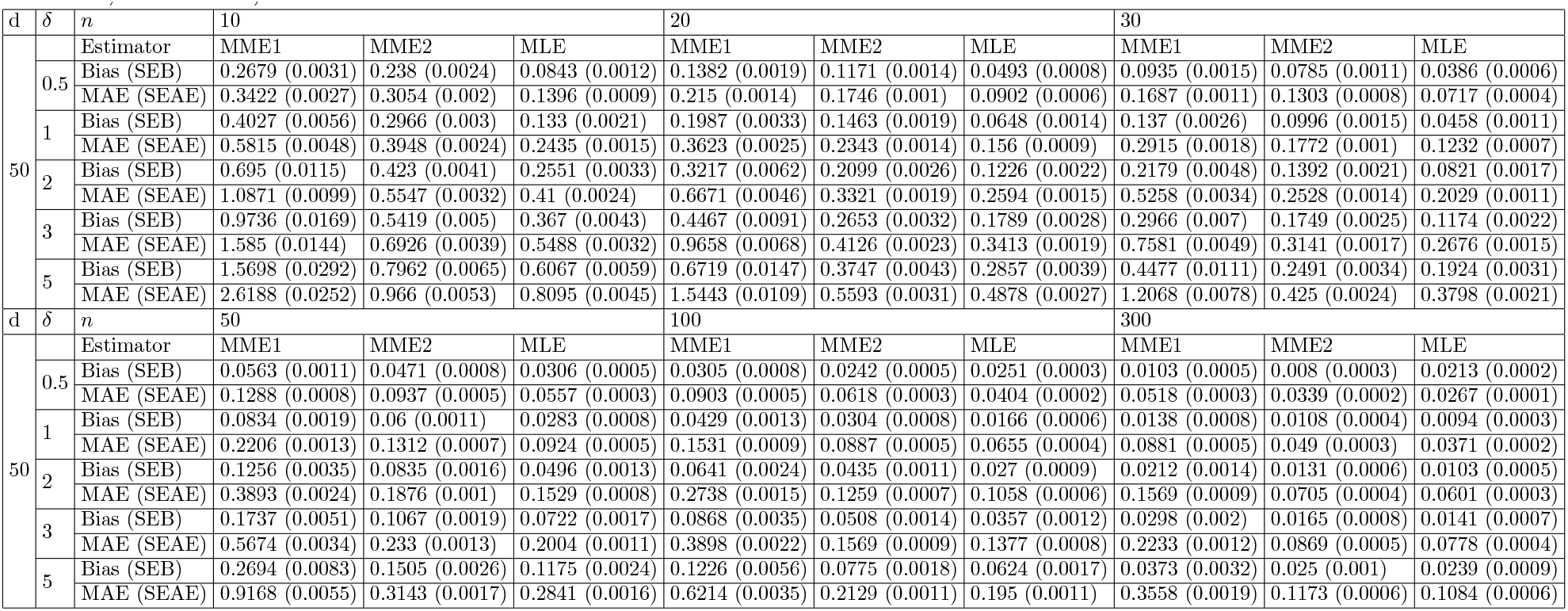
Large dimension estimation results for the symmetric Dirichlet with *δ* ∈ {0.5, 1, 2, 3, 5}, *d* = 50, *M* = 20, 000.

Remark 2.2: For the high dimensional (*d* = 50) case, the MLE outperforms both MME1 and MME2 in for all sample sizes in terms of SE, MAE, and SEAE. In terms of Bias, the MLE outperforms the other estimators for all but the largest sample size with the smallest parameter value, *n* = 300 and with common *δ* = 0.5. For small sample sizes (*n* = 10, *n* = 20, and *n* = 30) and common *δ* = 0.5, MME1 and MME2 take similar values with similar error ranges for BIAS and MAE while the MLE is much lower. For small sample sizes and common *δ* = 1, MME2 takes an intermediate position between the BIAS and MAE of MME1 and the MLE. For small sample sizes and common *δ* > 1, the Bias and MAE of MME2 grow closer to the Bias and MAE of the MLE as *δ* increases and as *n* increases. For large sample sizes (*n* = 10, *n* = 20, and *n* = 30), MME2 approaches but still underperforms compared to the MLE while MME1 has Bias, SE, MAE, and SEAE far above both. The exception is the case where *n* = 300 and common *δ* = 0.5, where MME1 and MME2 outperform the MLE and MME1 in terms of Bias (but not SE) and MME2 slightly outperforms MME1.

## 3 Asymptotic distribution of the MLE

Let *ψ* be the digamma function, the derivative of the natural logarithm of the gamma function. Let *ψ*^(1)^ be the trigamma function, the derivative of the digamma function. Additionally, define 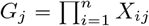 and G = (*G*_1_, *G*_2_, …, *G*_*d*_). For the Dirichlet distribution, the gradient vector of the likelihood, 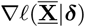, is

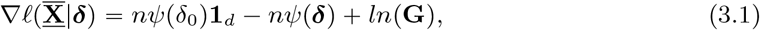

and the second derivative of the log-likelihood is

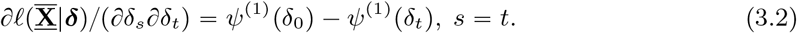

The second derivative is equal to 0 whenever *s ≠ t*. The semi-observed information is

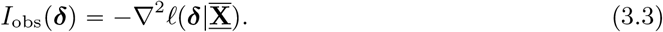

The Fisher information is then

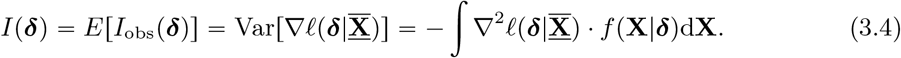

This holds given that the Fisher information exists and given regularity conditions guaranteeing commutativity of the integral and gradient operators with respect to *δ* (i.e. the probability density function is twice continuously differentiable and the support does not depend on the parameters).

For the Dirichlet distribution, the Hessian is

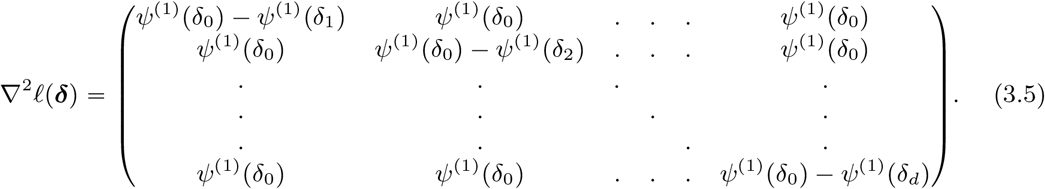

Since the Hessian is constant with respect to X,

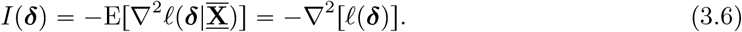

Given Equation 2.13, the dispersion matrix *S*_*e*_ for the bias of the MLE is

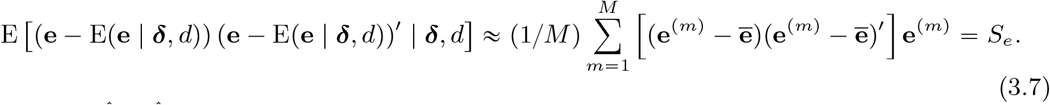

When 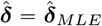, asymptotically the dispersion matrix *S*_*e*_ should converge to the asymptotic dispersion matrix Σ = (1/*n*)*I*^−1^(*δ*) as *n →* ∞. That is,

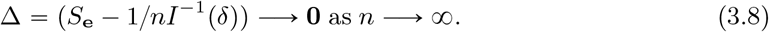

For the MLE of the Dirichlet paramter vector, the asymptotic dispersion matrix Σ can be expressed in closed form since the Fisher information is independent of the data. This means that there is no need to simulate the behavior of *S*_*e*_ since we can directly analyze Σ for various *δ*. The distribution of the inverse of Σ is shown in Figure 4. The diagonal elements represent the asymptotic variances (AVar) of 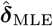 multiplied by *n*. For low values of dimension *d*, the diagonal elements are very similar to the nondiagonal elements, which represent the asymptotic covariances (ACov) of 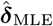 multiplied by *n*. For *d* = 50, the diagonal and nondiagonal elements diverge for higher values of *δ*. The variances and covariances also increase for higher values of *δ* compared to lower ones. Figure 5 shows the values of the asymptotic dispersion matrix as well as the asymptotic correlations for elements of the MLE for the Dirichlet distribution with dimension *d* = 2. For examples of the *d* = 3 case, see Figures 12, 13, and 14 in the Appendix.

**Figure 3.**
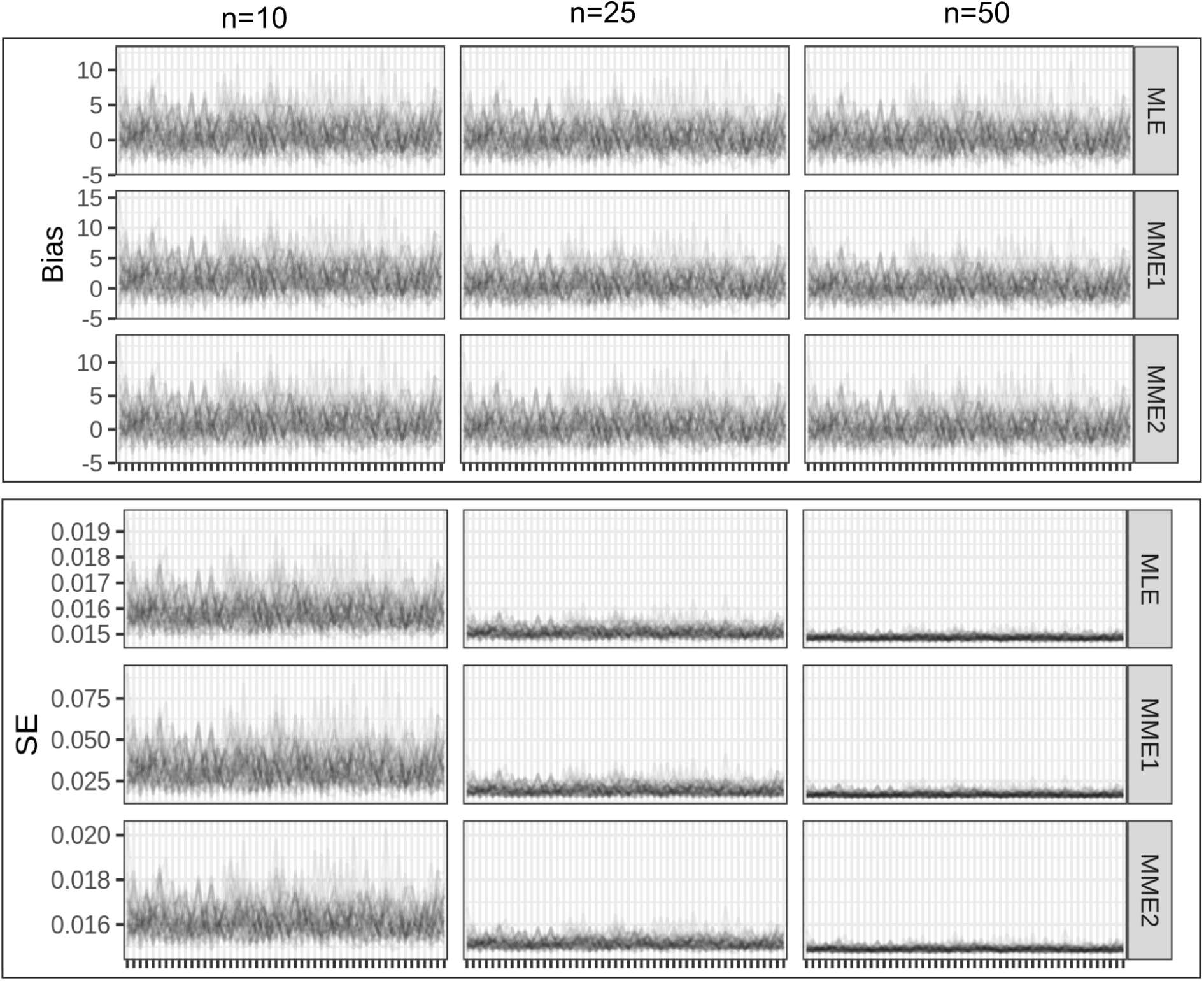
Estimation results for the asymmetric Dirichlet distribution with *d =* 50. Bias for (**a**) and SE (**b**) for 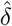 are plotted by *δ*_*i*_ for all three estimators for sample sizes of 10, 25, and 50. Parameter values are iid Gamma(5, 1). The number of simulations per iteration, *M*, is 20,000. The number of iterations is 50.

**Figure 4.**
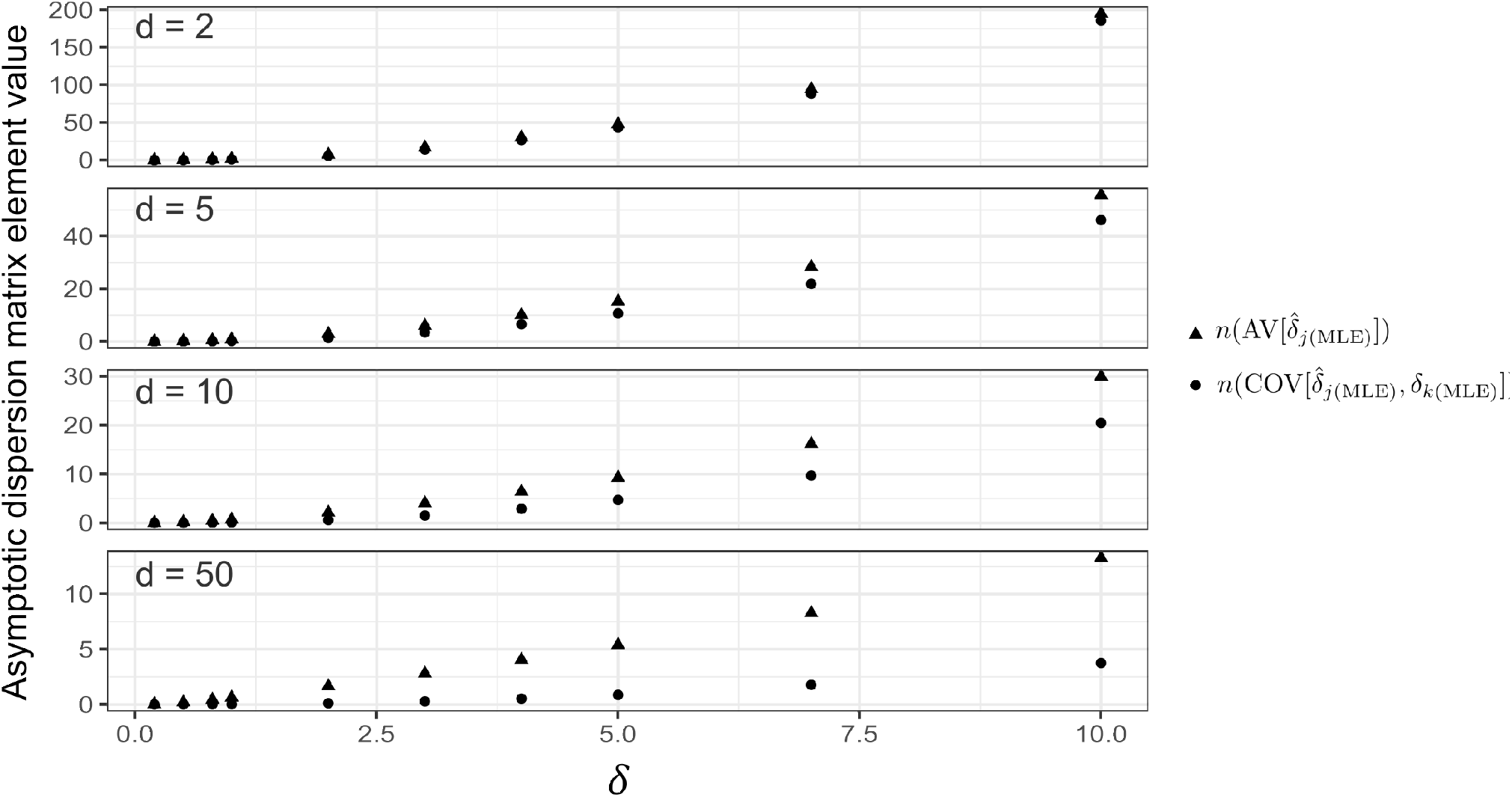
Diagonal and nondiagonal elements of the asymptotic dispersion matrix for the symmetric Dirichlet distribution for *d* = 2, 5, 10, 50.

**Figure 5.**
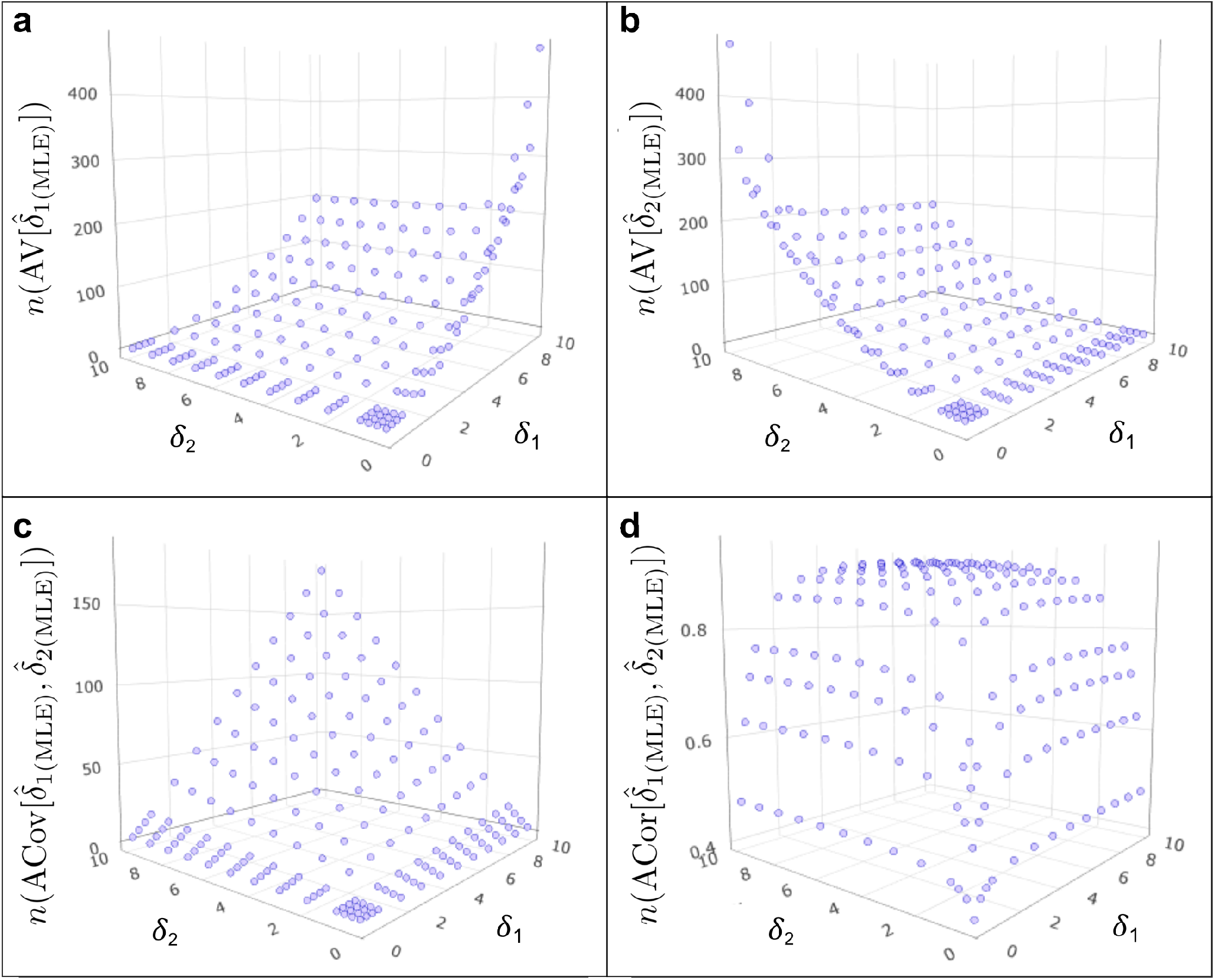
Asymptotic dispersion matrix element values for the MLE of the Dirichlet distribution with dimension *d* = 2. The diagonal elements of the asymptotic dispersion matrix (**a, b**) represent the asymptotic variances of 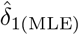 and 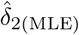. The nondiagonal elements shown in **c** represent the asymptotic covariances. The asymptotic correlations are shown in **d**.

In Figures 4 and 5, the asymptotic covariance and correlation values appear to always be positive for different choices of *δ*. Indeed, this must always be the case as we will now prove. Let us first rewrite the Fisher information, which is here the negation of the Hessian, as the sum of two matrices in the manner of Ng, Tian, and Tang (2013). [21]

Define G as

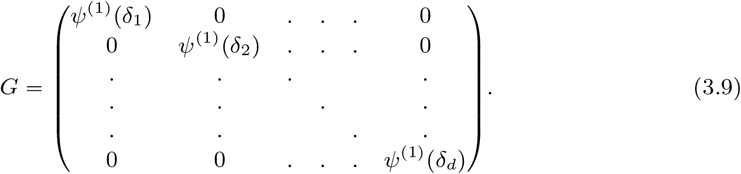

Let 1_*d*_ be the standard d-length unit vector. Then the Fisher information is

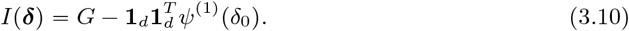

In his 1981 paper[30] Kenneth S. Miller showed that given two matrices *A* and *B*, if *A* and *A* +*B* are invertible and *B* is of rank 1, then the following holds:

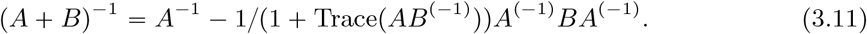

For our case, *G* and 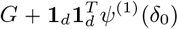 are positive semi-definite and 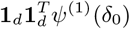 can be trivially decomposed into the product of two vectors and so must be of rank

1. With these conditions satisfied, we use Equation 3.10 to rewrite Σ as

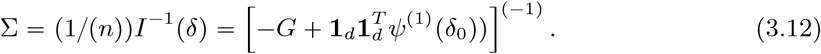

Given the above formulation of Σ and using Equation 3.11, we have that every element of Σ is positive if for all *j, k*,

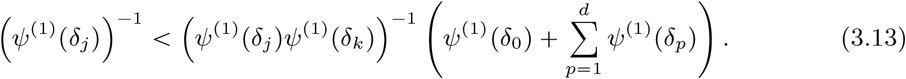

This is true if and only if

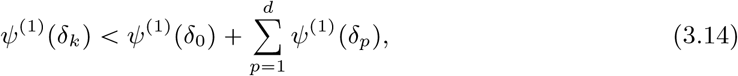

which implies that

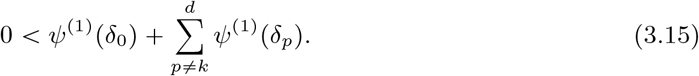

The trigamma function *ψ*^(1)^ is monotonic decreasing and approaches but never reaches 0 as *δ →* 0, so Equation (3.15) is always true. Therefore, the asymptotic covariances (and thus the asymptotic correlations) between the elements of the MLE of the Dirichlet parameter vector are always positive. For large sample sizes, elements of the Dirichlet MLE will always be positively correlated with each other. This correlation of the elements of the MLE is in contrast to the correlation of the elements of a Dirichlet distributed random vector, which is strictly negative as shown in Equation 2.9.

## 4 Application on real-life microbiome data

### 4.1 Existing GoF tests for the Dirichlet distribution

When working with real world data, an important first step to verify the applicability of a particular distribution is GoF testing. Although GoF tests for the Dirichlet distribution have been proposed, they have not yet been applied to evaluate the distribution’s fitness for application on different populations of microbiome data.[24] Previous work on GoF tests for the Dirichlet distribution has formulated three different tests: the Distance Covariance Test (DCT), the Dirichlet Energy Test (DET), and the Triangle Test (TRT).[24] The DCT applied to Dirichlet distribution builds on previous work done by Connor and Mosimann (1969), Daroch and Ratcliff (1971), Fabius (1973), and James and Mosimann (1980) on the Dirichlet distribution and independence.[31, 32, 33, 34] The TRT was created by Bartoszyński, Pearl, and Lawrence (1997) by utilizing universal geometric properties of data and distributions.[35] Unfortunately, the DCT, DET and the TRT are all impractical for high dimensional microbiome data. The formulation of a proper GoF method for high dimensional Dirichlet distributions remains an open problem.

### 4.2 Datasets and results

We apply our estimation method to four different datasets: a small sample gut microbiome dataset from Lahti et al. (2013), a large sample gut microbiome dataset from Lahti et al. (2014), a small sample soil microbiome dataset from Zhou et al. (2011), and a small sample soil microbiome dataset from Caporaso (2011).[36, 37, 38, 39] The datasets will be referred to henceforth as Gut1, Gut2, Soil1, and Soil2, respectively. The Gut1 dataset contains 22 samples of 130 different taxa. The Gut2 dataset contains 1006 samples of 130 different taxa. A heatmap of Gut2 is shown in the Appendix as Figure 14. The Soil1 dataset contains 14 samples of 16,825 taxa. The Soil2 dataset contains 3 samples of 19,216 taxa. Due to the extremely large *d* values resulting from the soil datasets, we followed the cleaning procedure of Aubert, Schbath, and Robin (2021) and removed all taxa that did not represent at least 0.5% of at least one sample.[40] This brought the number of taxa in Soil1 to 104 and the number of taxa in Soil2 to 62. Using a sample dataset of *d* = 2 created by agglomerating the first and second 65 taxa of Gut1, the DCT GoF test gives p-values of 0.3762 and 0.3762, indicating that we fail to reject the null hypothesis that the data is distributed according to a Dirichlet distribution. Because the dimension of the full datasets renders the three previously discussed GoF tests unusable, we proceed under the assumption that these datasets are Dirichlet distributed.

We compare the results of our model to those of the Dirichlet-Multinomial Method (DMM) as used in Holmes, Harris, and Quince (2012).[23] This method is summarized in the Appendix (Section 8.3). The results for estimates of both models are shown in Figure 6, Figure 7, Figure 8, and Figure 9. Since comparing the raw values of the Dirichlet parameters is not informative, we compare the estimates of the expected relative abundances (Equation 4.3, which is obtained from Equation 2.5).

**Figure 6.**
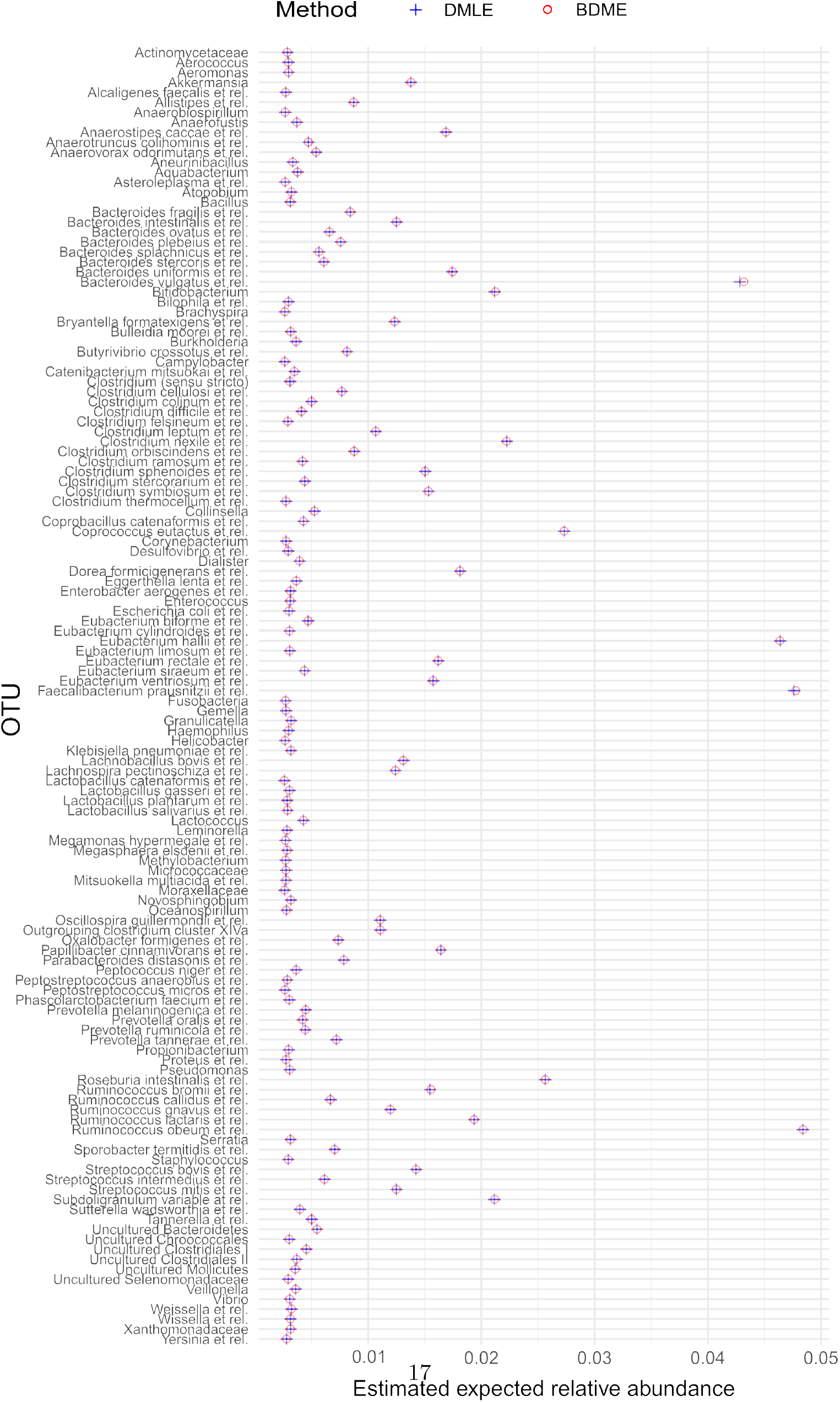
Estimated expected relative abundance of gut microbiome data (Gut1) using both the Dirichlet Maximum Likelihood Estimator (DMLE) and the Bayesian Dirichlet Multinomial Estimator (BDME) derived from the model described in Holmes, Harris, and Quince (2011).[23] This dataset is from Lahti *et* al (2014).[36] The sample size *n* = 22 and the dimension *d*=130.

**Figure 7.**
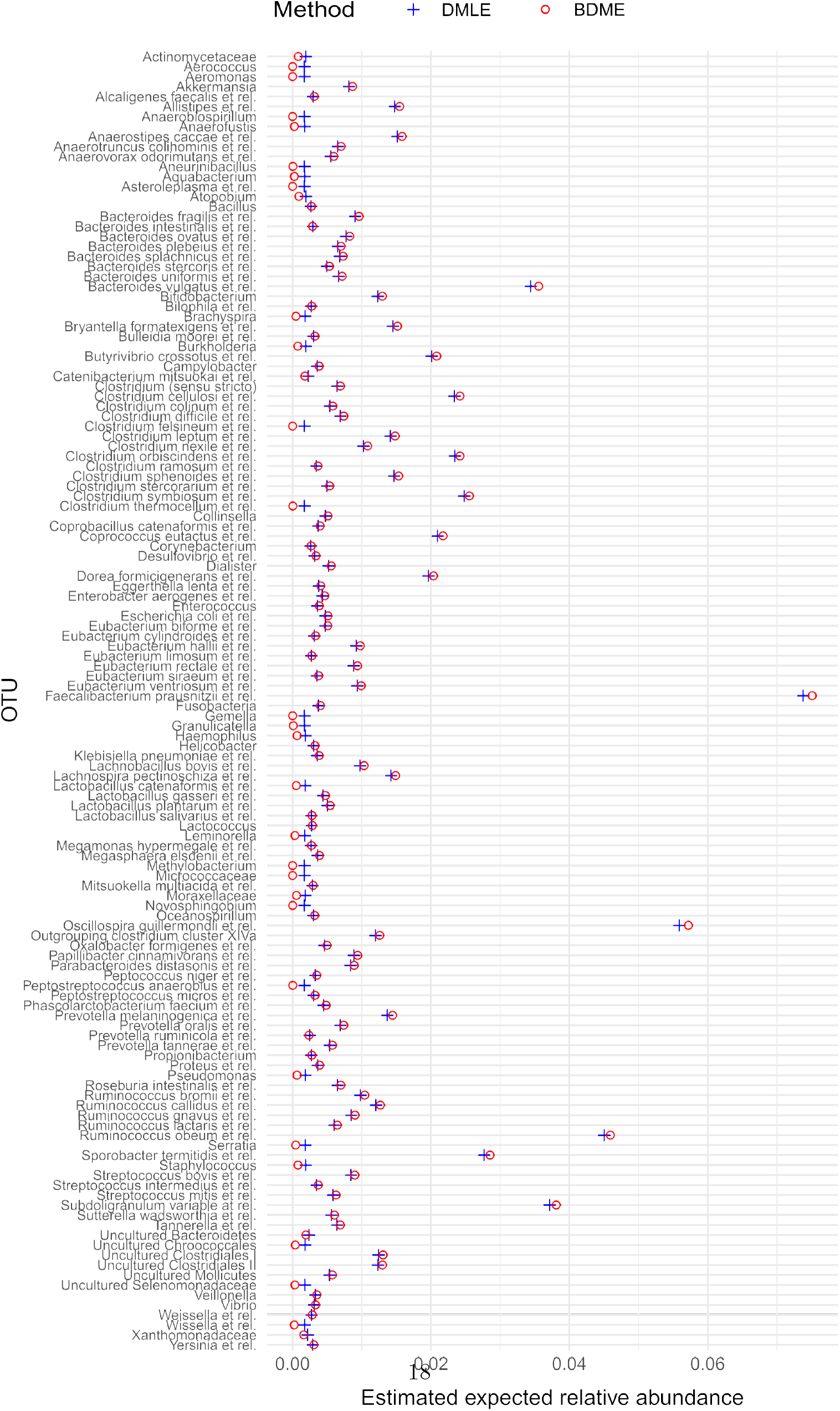
Estimated expected relative abundance of gut microbiome data (Gut2) using both the Dirichlet Maximum Likelihood Estimator (DMLE) and the Bayesian Dirichlet Multinomial Estimator (BDME) derived from the model described in Holmes, Harris, and Quince (2011).[23] This dataset is from Lahti *et* al (2014).[37] The sample size *n* =1006 and the dimension *d* = 130.

**Figure 8.**
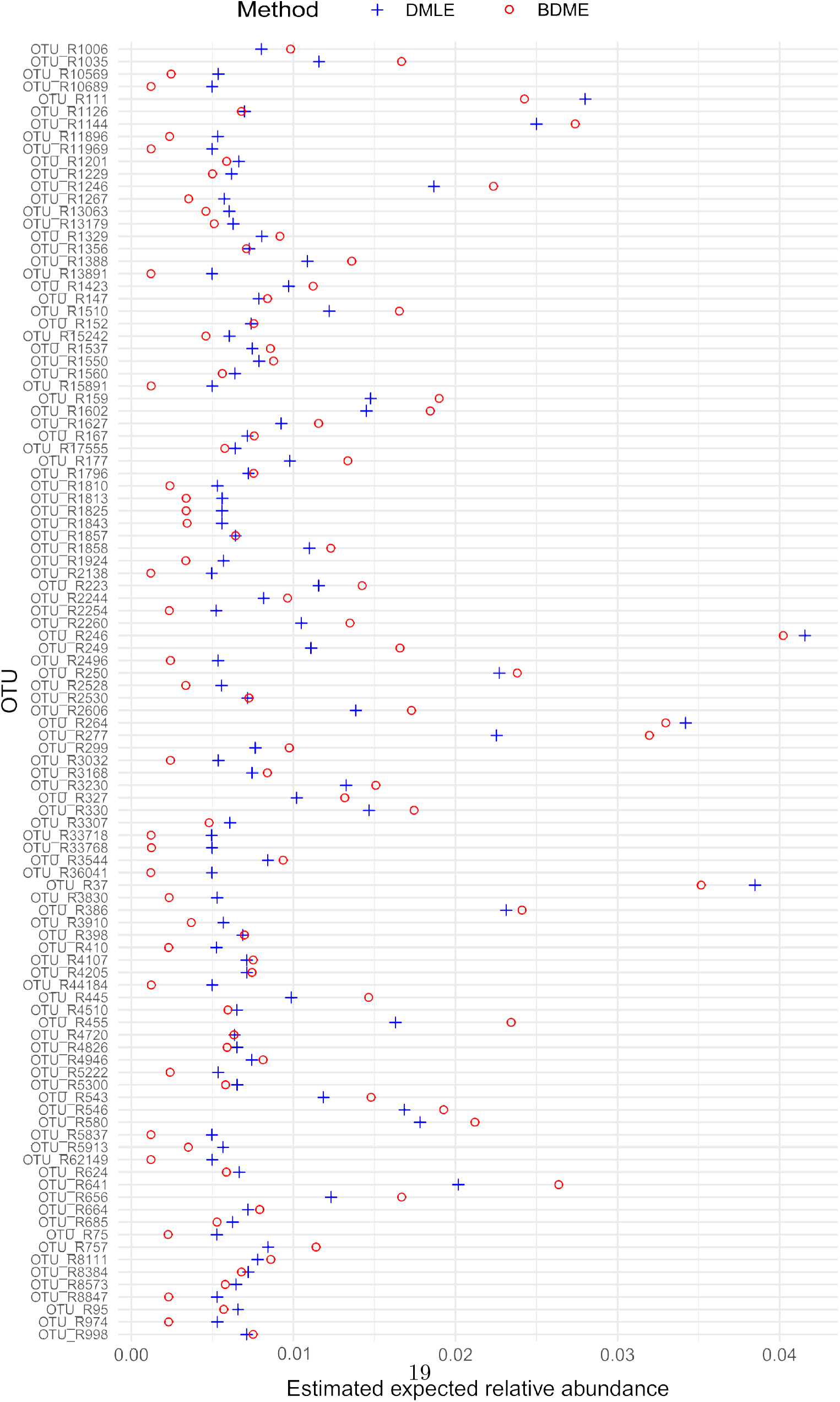
Estimated expected relative abundance of soil microbiome data (Soil1) using both the Dirichlet Maximum Likelihood Estimator (DMLE) and the Bayesian Dirichlet Multinomial Estimator (BDME) derived from the model described in Holmes, Harris, and Quince (2011).[23] This dataset is from Zhou *et* al. (2011).[38] The sample size *n* = 14 and the dimension *d* = 104.

**Figure 9.**
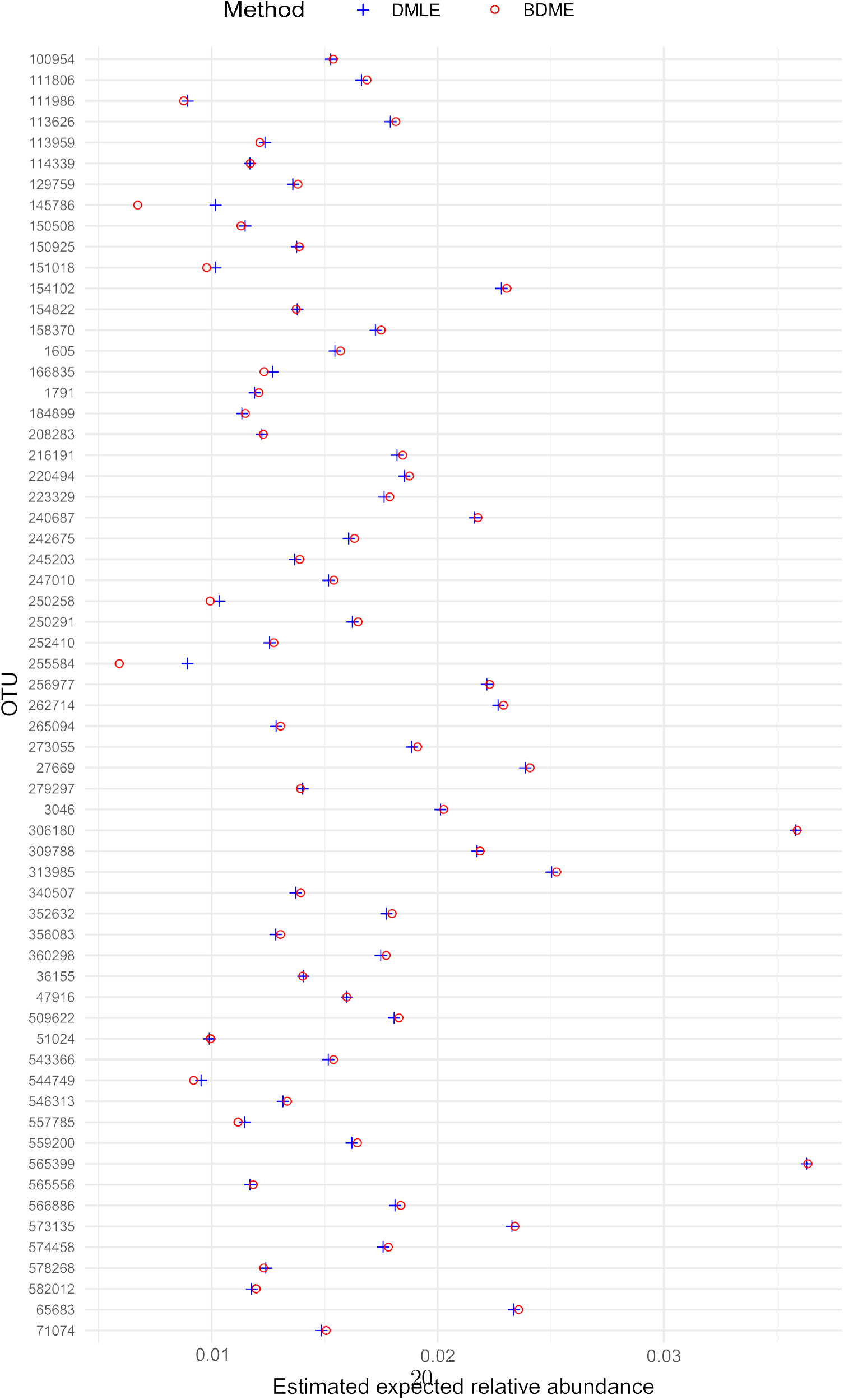
Estimated expected relative abundance of soil microbiome data (Soil2) using both the Dirichlet Maximum Likelihood Estimator (DMLE) and the Bayesian Dirichlet Multinomial Estimator (BDME) derived from the model described in Holmes, Harris, and Quince (2011).[23]This dataset is a subset of the samples from the well-studied GlobalPatterns dataset.[39] The sample size *n* =3 and the dimension *d* = 62.

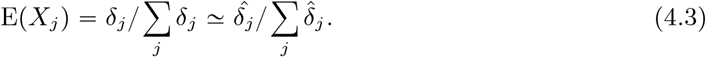

### 4.3 Remarks

In each case, estimates from both methods were similar but the BDME took a factor of 5 to 10 more time to complete. Estimates for both models appear much closer to each other for the Gut microbiome datasets (Figures 6 and 7) than the Soil datasets (Figures 8 and 9). This is in part due to the nonexistant sparsity of the gut microbiome datasets and larger samples sizes. Both gut microbiome datasets sequenced with microarray technology, which has lower accuracy at low count levels. It also works by tagging a preselected number of specific taxa, which is why *d* = 130 for both gut microbiome datasets. Soil1 was sequenced with pyrosequencing while Soil2 was sequenced with the more advanced but related Illumina sequencing. The differences in both parameter value and estimated expected relative abundance for Gut1 are shown in Table 4. The BDME estimates for 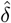 are always higher than the DMLE ones, but the estimators give similar outputs for estimated expected relative abundance, which is the more useful and informative measure in practice. As described in Lahti et al., many microbiomes exhibit bimodal distributions of relative abundances with many taxa being relatively sparsely observed and a few taxa dominating in most samples. This trend is reflected in Figure 6 with the majority of taxa occupying a low relative abundance region E(*X*_*j*_) < 0.01 and only a few others occupying a region of high relative abundance E(*X*_*j*_) > 0.01. In contrast to previous analysis of this data, however, E(*X*_*j*_) provides a summary of the relative abundance for taxa across samples as opposed to a per-sample bimodal distribution of relative abundances.

**Table 4:**
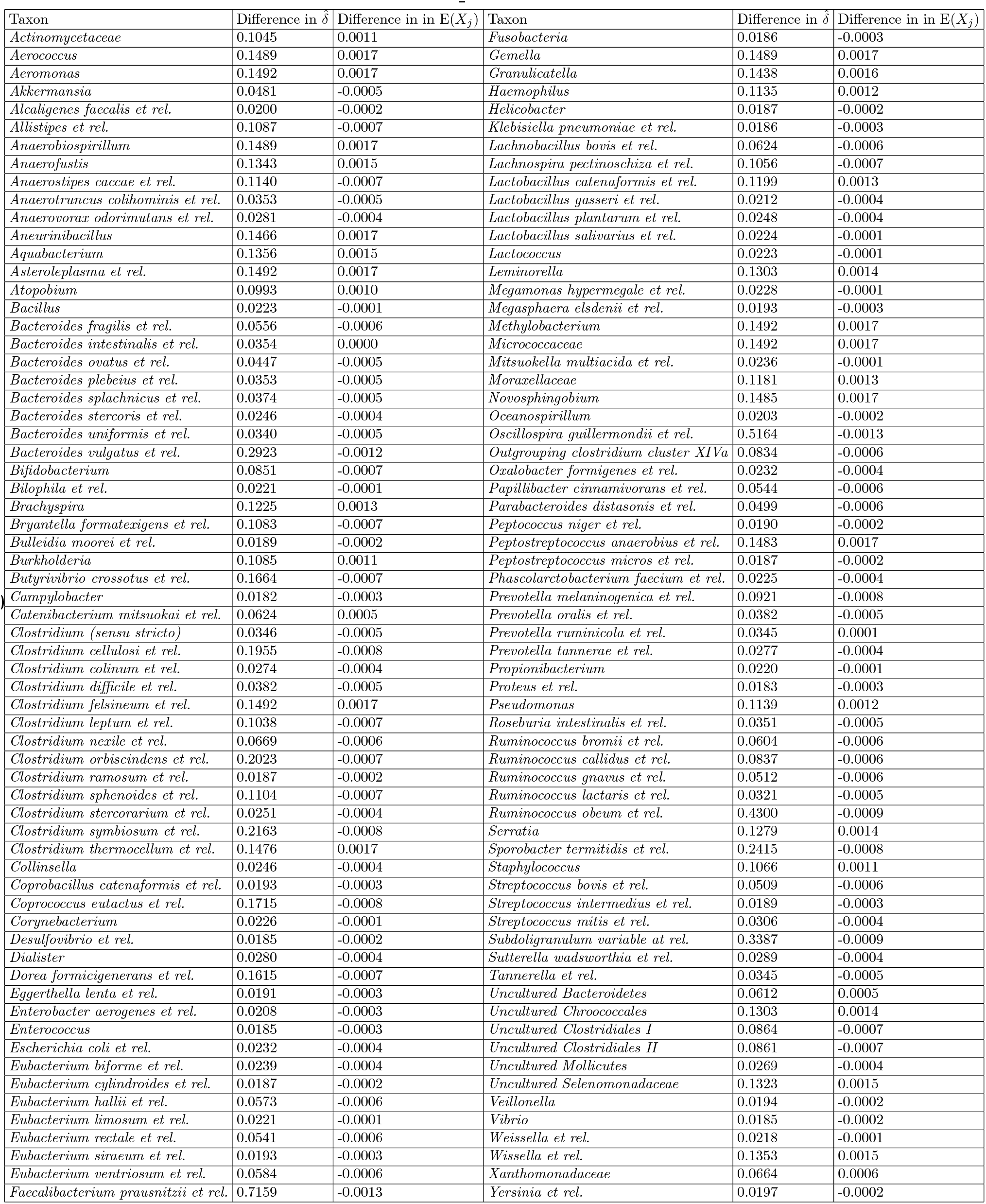
Comparisons between the parameter estimates of the DMLE and the BDME derived from the Bayesian Dirichlet-Multinomial model described in Holmes, Harris, and Quince [23]. The dataset used was Gut1 and differences are calculated with respect to the DMLE.

**Table 5:**
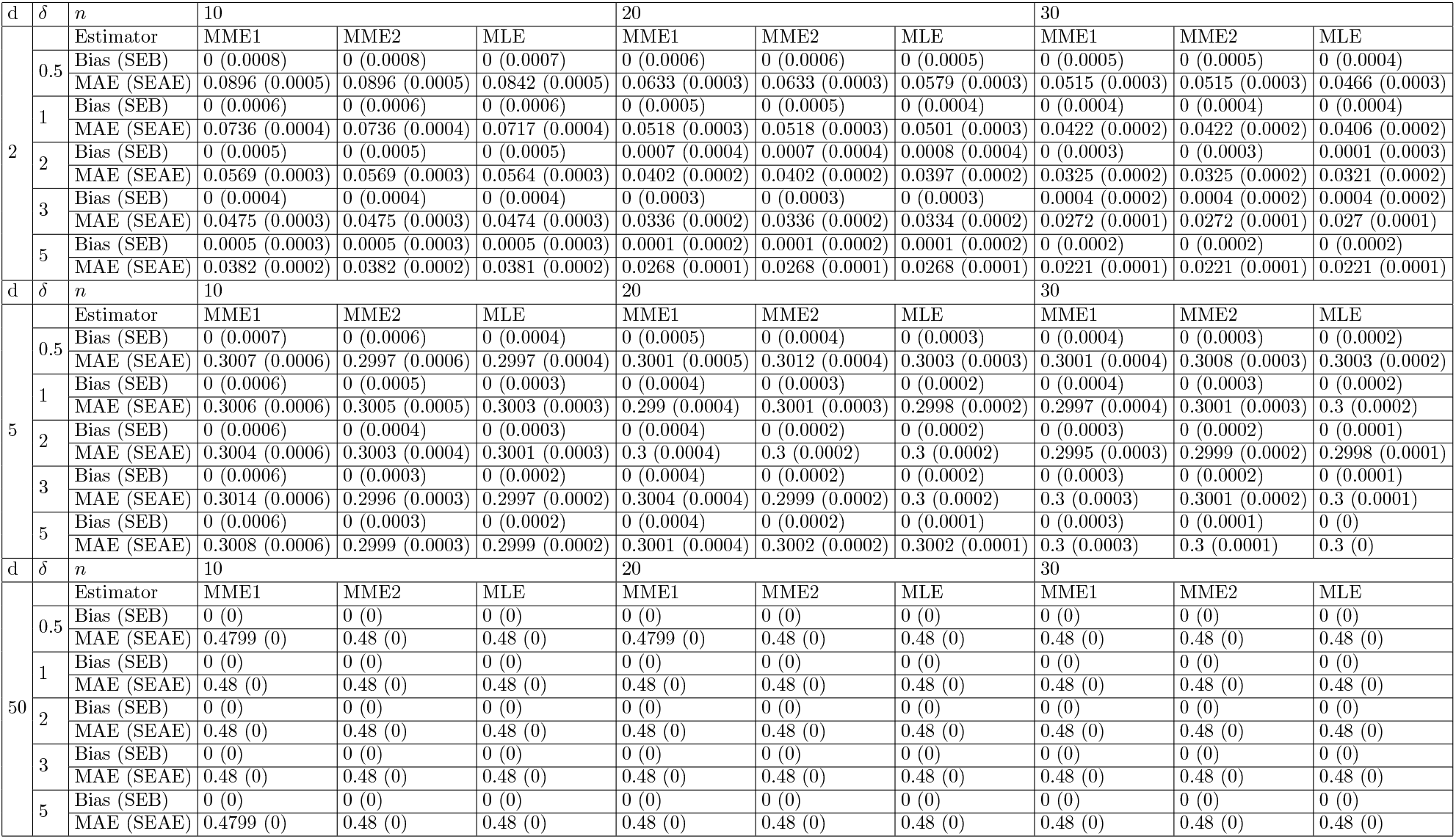
Small sample expected relative abundance estimation results for the symmetric Dirichlet with *δ* ∈ {0.5, 1, 2, 3, 5}, *d* ∈ {2, 5, 50}, *M* = 20, 000.

**Table 6:**
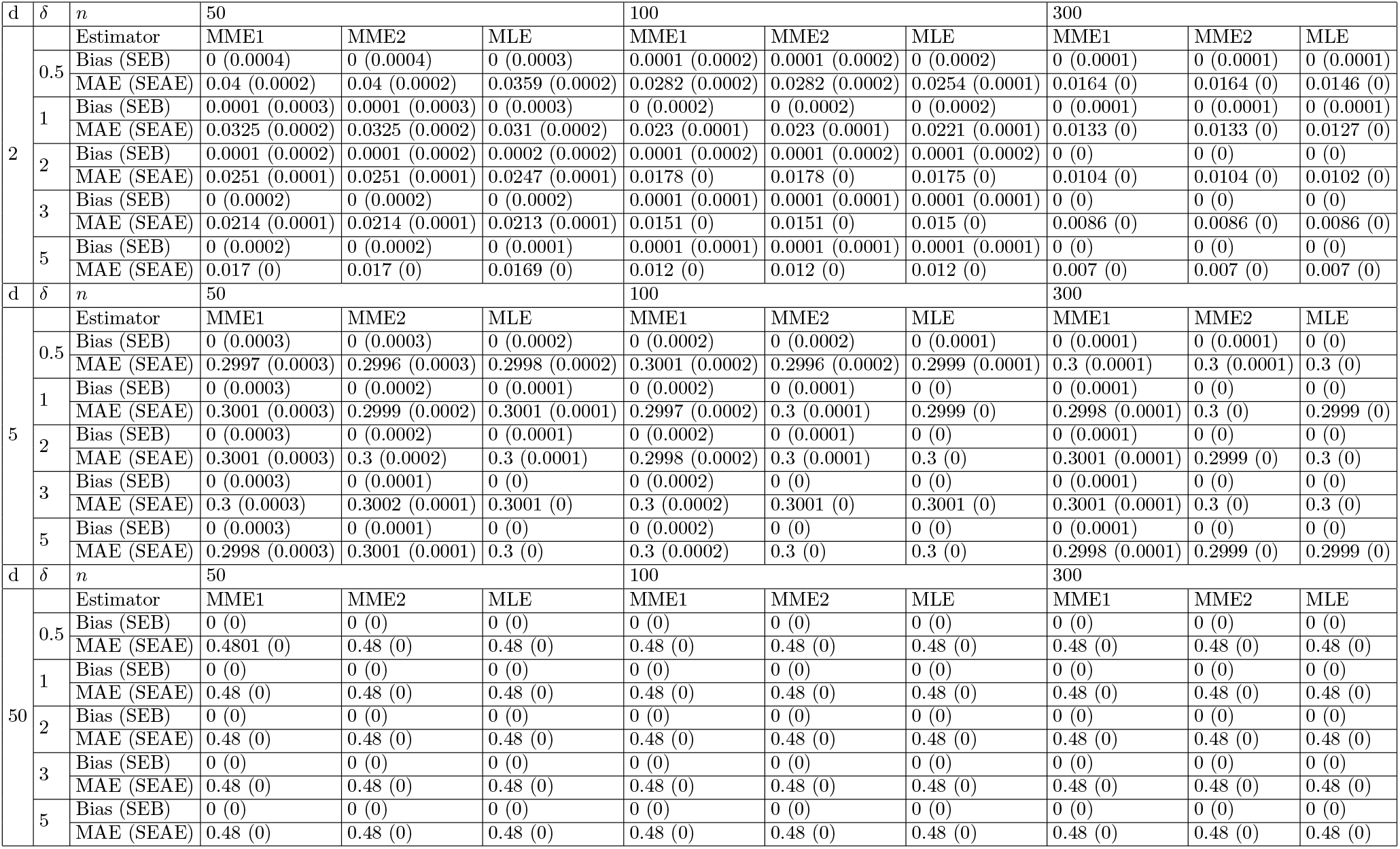
Large sample expected relative abundance estimation results for the symmetric Dirichlet with *δ* ∈ {0.5, 1, 2, 3, 5}, *d* ∈ {2, 5, 50}, *M* = 20, 000.

The differences in estimated expected relative abundance are more stark between the Soil1 and Soil2. Soil1 (Figure 8) shows a large difference between the two estimators while Soil2 (Figure 9) shows almost no difference. This is despite the fact that Soil2 has a much smaller sample size (*n* = 3) compared to Soil1 (*n* = 14).The reason for this difference is sparsity. Soil1 is 54% sparse and Soil2 is only 1% sparse post cleaning, but pre-cleaning Soil1 is 93% sparse and Soil2 is 65% sparse. So while Soil1 is an inherently more sparse dataset, the low total counts also affects the cleaning procedure since singletons represent much larger percentages of the library size and the cleaning procedure is percentage based. This means that the overall structure of Soil1 is much different than Soil2, leading to the difference in estimation consistency. With the large amount of zeroes present in the dataset, the DMLE pulls most of its estimations towards the interior of the parameter space in comparison to the BDME. This also highlights a potential weakness of the Soil1 dataset, which is that the low total counts indicate that proper sequencing depth may not have been achieved, meaning that the sample may not be properly representative of the population from which it was taken.

The bias and SE of the BDME could not be quantified in the simulation study along with the other estimators because the BDME is calculated off of raw counts only, which cannot be produced by the methodology utilized for our simulations. In order to compare the efficacy of the BDME and the DMLE, we used nonparametric bootstrapping and compared the bias and RMSE of the estimators. This was only performed on Gut2 because it was the only dataset with a large enough sample size for nonparametric bootstrapping to give trustworthy results. The number of bootstrap replications performed was *k* = 10, 000. Because of the possibility of having all-zero samples during resampling, the boostrapping was performed after shifting all of the raw counts up by 1, meaning that taxa that were previously recorded as absent would have a count of 1 instead. This is a commonplace strategy for dealing with zero-inflated data. See, for example, Huang and Peddada. (2020)[20] The comparison results are shown in Figure 10. The estimators have opposite biases. That is, when BDME is positively biased, DMLE is negatively biased, but the opposite also holds true for most cases. In terms of magnitude, however, the BDME has a lower bias than the DMLE. The value of both biases is extremely small, however, as shown when comparing the scales of Figure 10 to Figure 7. Both estimators had very similar similar RMSE.

**Figure 10.**
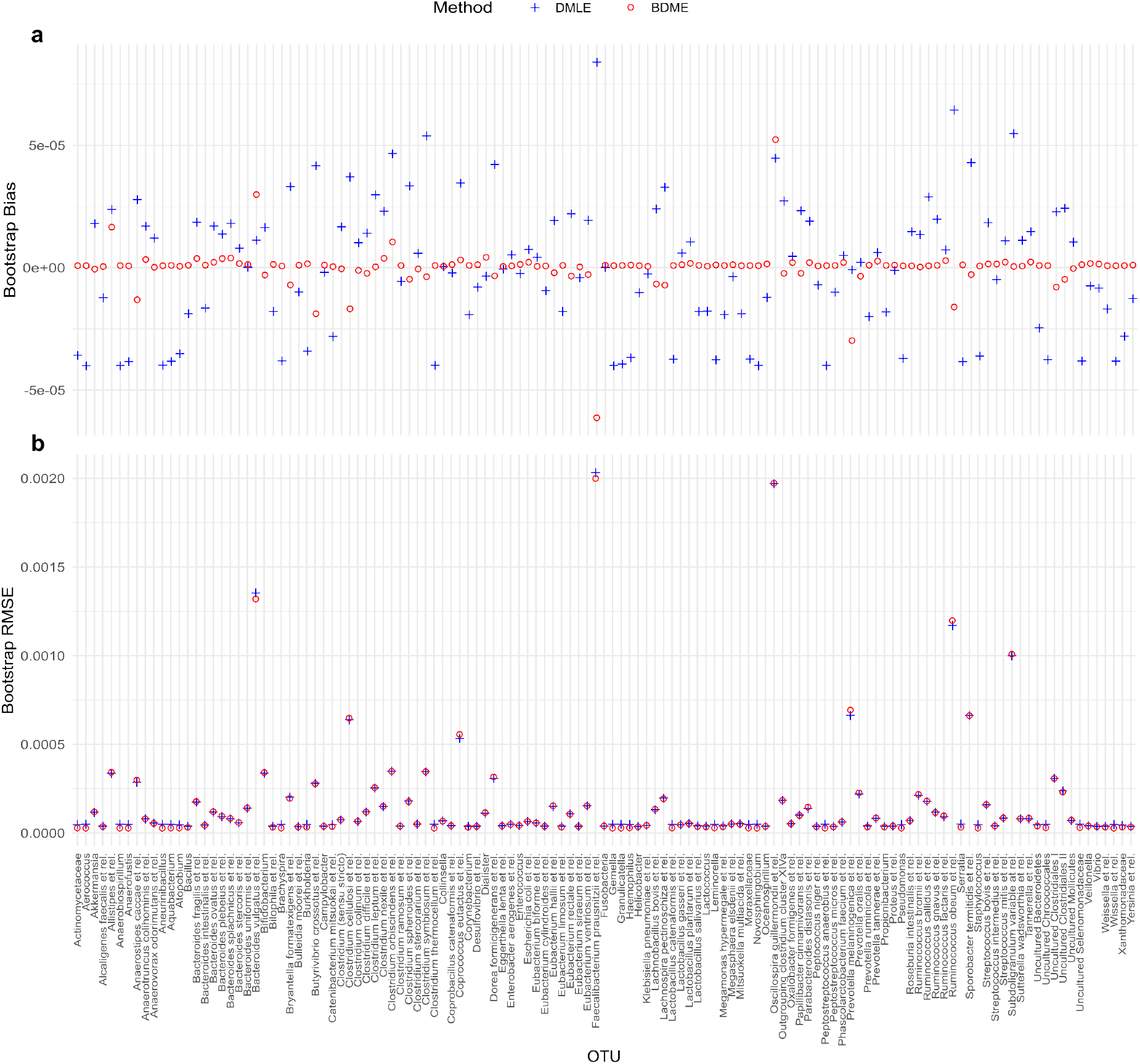
Comparison of estimator bootstrap bias (**a**) and RMSE (**b**) between BDME and DMLE using nonparametric bootstrapping. The number of replications was *k* = 10, 000.

**Figure 11.**
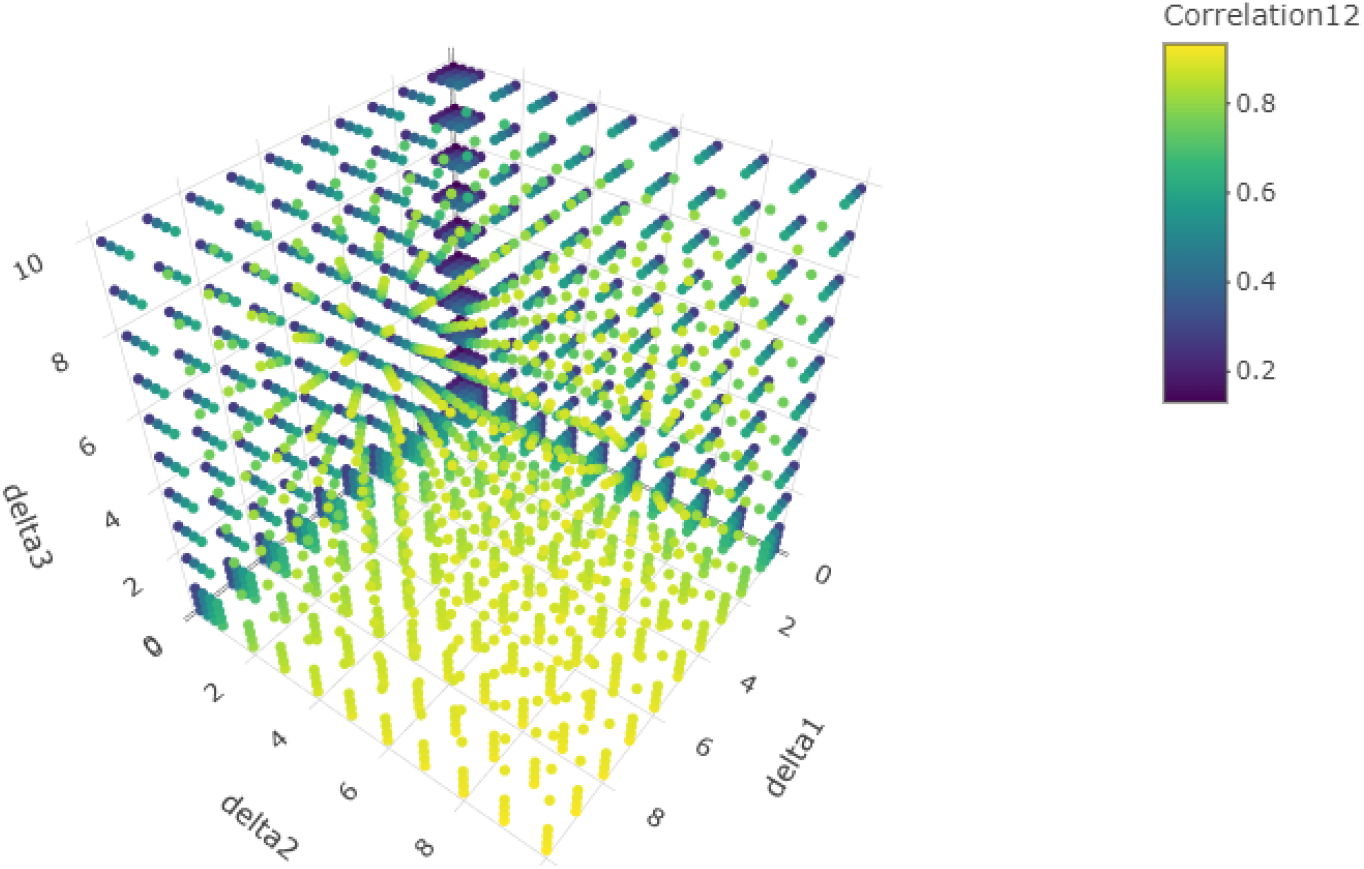
Assymptotic correlation between the first and second component of the MLE of the symmetric Dirichlet distribution with *d* = 3.

**Figure 12.**
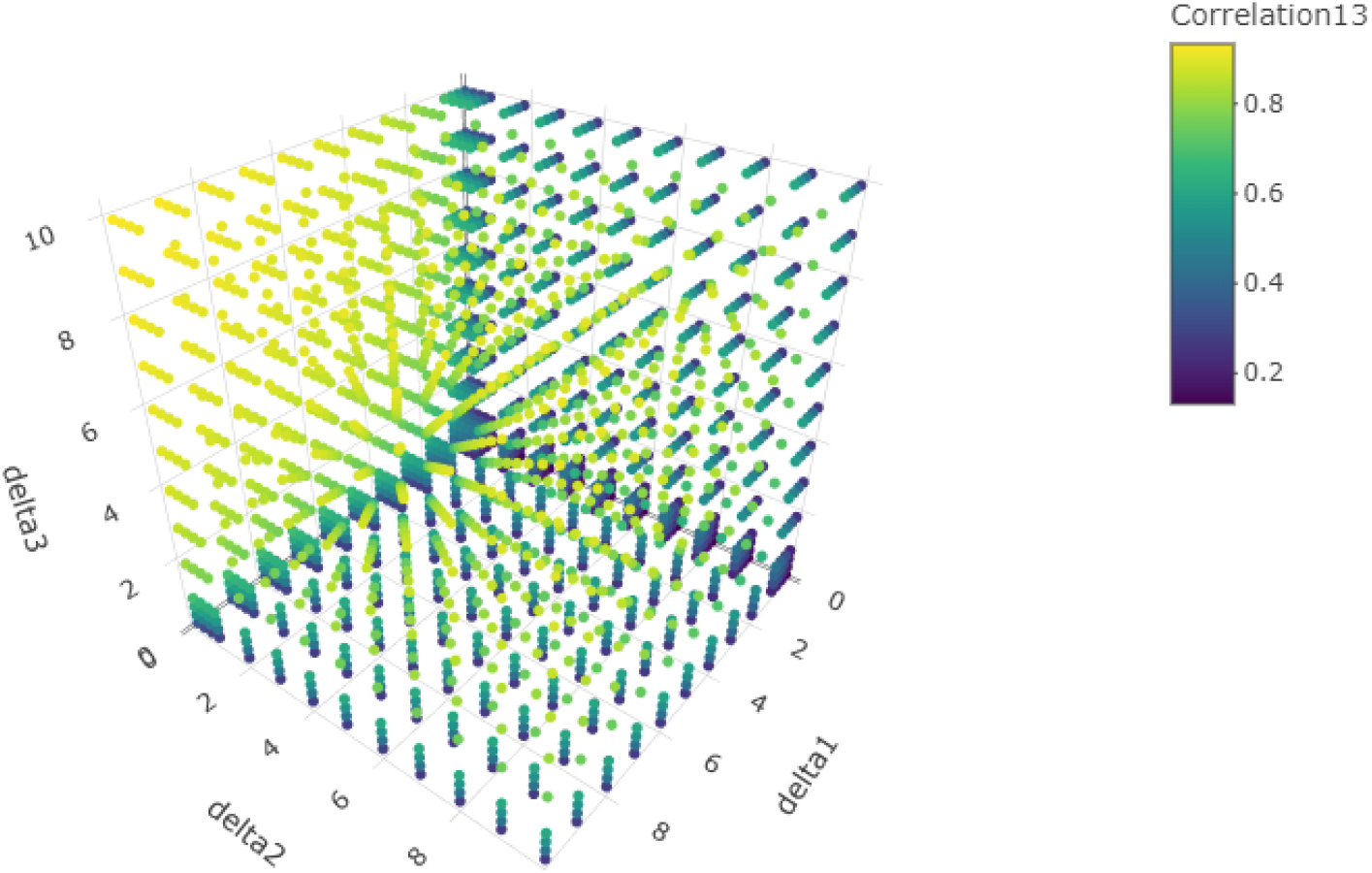
Assymptotic correlation between the first and third component of the MLE of the symmetric Dirichlet distribution with *d* = 3.

**Figure 13.**
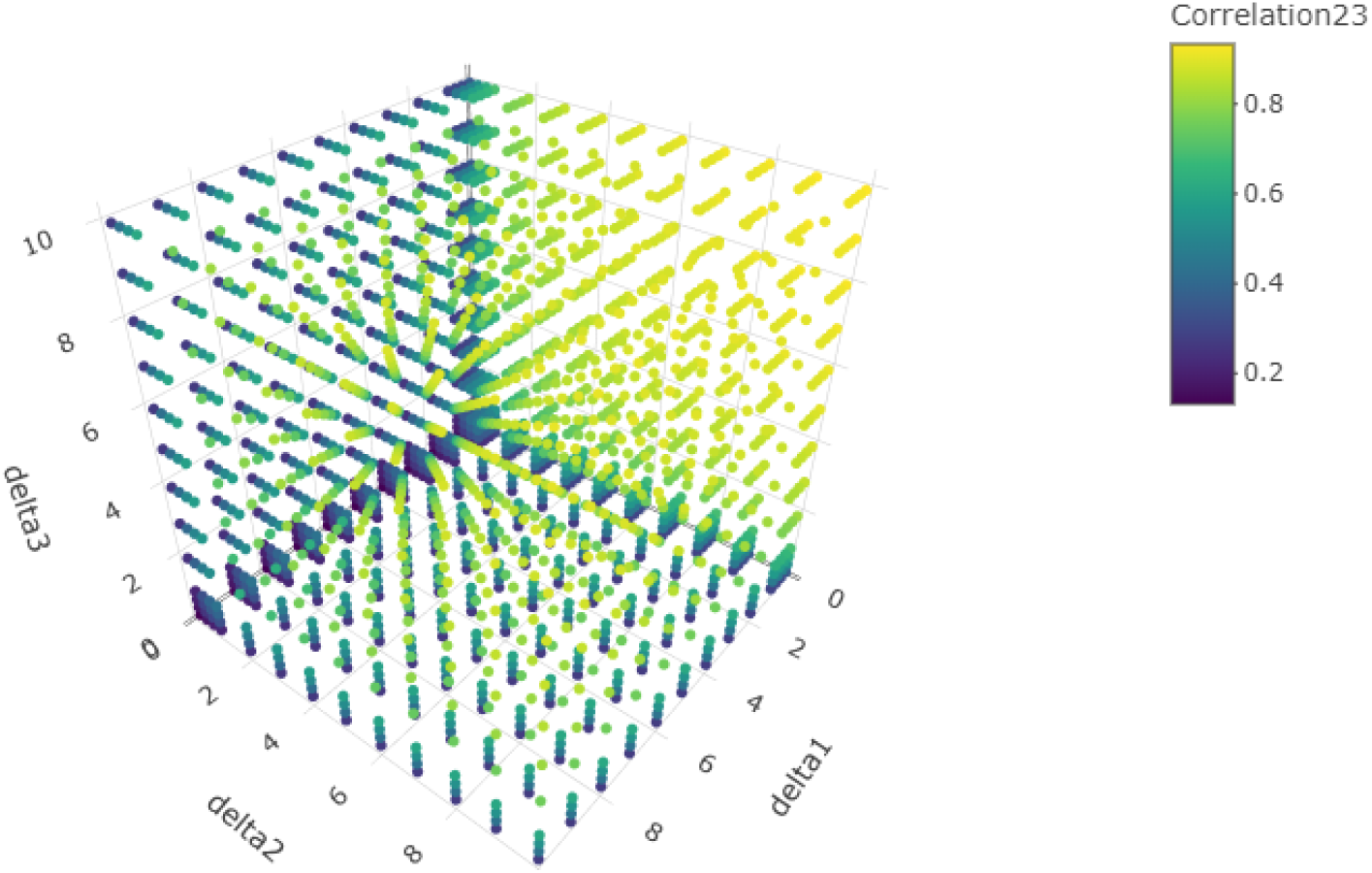
Assymptotic correlation between the second and third component of the MLE of the symmetric Dirichlet distribution with *d* = 3.

**Figure 14.**
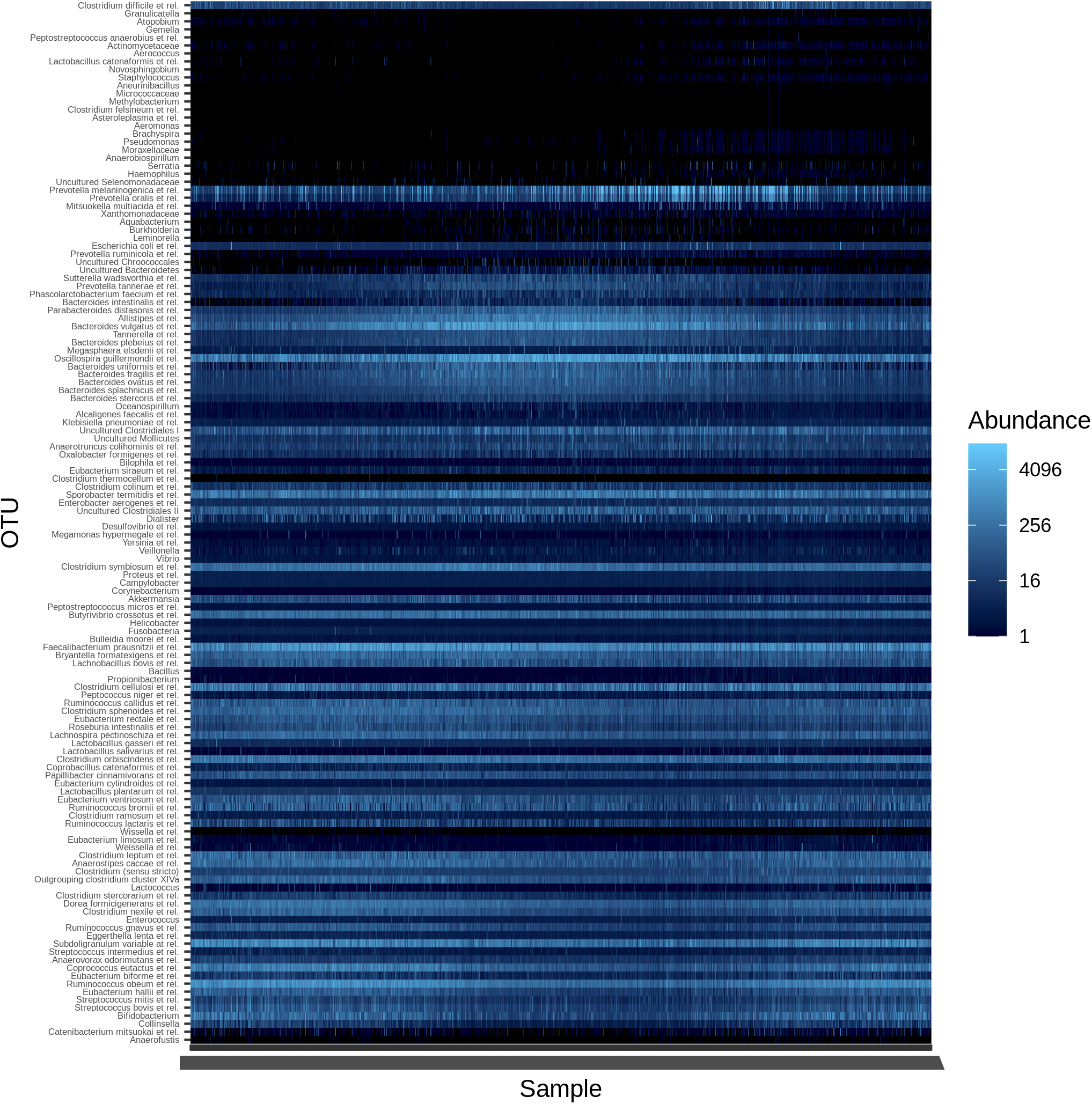
Heatmap of the taxonomic absolute abundances of gut microbiota from the dataset in Lahti et al (2014) [37].

## 5 Concluding remarks

In this study, we improved upon existing literature in Dirichlet parameter estimation and compared the performances of the MLE as well as two different MMEs for low and high sample sizes. While the MLE is expected to outperform the MMEs for large samples, we found that 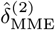 exhibited similar levels of bias and MAE across all sample sizes and values of dimension *d* and parameter vector ***δ***. The asymptotic variances and covariances of the dataset were identified via the asymptotic dispersion matrix. Although some work in Dirichlet parameter estimation has been done before, this is the first work to perform a comprehensive study comparing the bias and MAE of these estimators. This is also the first study to look at high dimensional estimation of the Dirichlet parameter vector, which is critical to consider for microbiome analyses.

### 5.1 Statistical findings

Parameter estimation for the symmetric case of the Dirichlet distribution should become least biased as the dimension *d* increases since there is technically only a single parameter value (the common *δ*) being estimated while there is an increase in information. This expectation was born out in simulation for all symmetric cases except for when *δ* < 1. For this case, the bias of the MLE increased as *d* increased for constant *δ* and sample size *n*. This trend held for the asymmetric cases as well. When any *δ*_*j*_ < 1, the bias for the MLE of that *δ*_*j*_ was higher for high *d* than for low and for cases similar *d* with no *δ*_*j*_ < 1. For future work involving statistical tests of the Dirichlet distribution, these results mean that using the MLE could be problematic for cases where true common *δ* < 1 or where any particular *δ*_*j*_ < 1. Increased bias for an estimator could result in an inflated empirical size for a test for the value of *δ*, for instance. This is not simply a problem for simulated distributions, either. The parameter estimations performed in Section 4.2 found that many sparsely represented taxa have 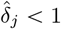.

Overall, the performance of the MLE was superior to MME1 and MME2 in terms of both bias and MSE for the majority of cases. To obtain a better understanding, we investigated the asymptotic behavior of the MLE using the asymptotic dispersion matrix Σ and proved that the asymptotic covariance (and therefore the asymptotic correlation) between elements of 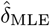 is always positive. This finding is of note because the correlation of Dirichlet distribution parameters is strictly negative. This positive correlation between estimators could help explain the increased bias for the MLE under certain conditions. This is also important in the context that there are examples of microbiome count data exhibiting a mixture of positive and negative Pearson correlations, which has been used to criticize the potential choice of the Dirichlet distribution as a model for microbiome data in the work of Mandal et al. (2015) and others.[20]

### 5.2 Relevance to microbiome research

It is important to have estimation of the relative abundance in the population when analyzing microbiomes, without which it is difficult to obtain any viable information. In this regard, the Dirichlet MLE does that comparably well to the Dirichlet-Multinomial algorithm while using considerably fewer computational resources, especially if the estimation procedure is parallelized. For the MLE, sample sizes of 30 and beyond are very consistent in terms of low bias and MSE for even very large dimensions. For ANCOM-BC, the recommended minimum sample size is 10, under which the false discovery rate has been shown to increase past acceptable levels.[10] For applications where this sample size is not achievable, parametric bootstrapping may serve to improve the performance of statistical tests based on the MLE. In the future, Dirichlet modeling could allow for direct implementation of statistical tests for microbiome populations.

Goodness of fit testing is the standard practice by which the the applicability of the Dirichlet distribution for microbiome data could be confirmed. Unfortunately, current GoF tests that have been developed for the Dirichlet distribution are unsuitable for microbiome data because of the very high dimension and the frequency of zero values. Additionally, these tests have not been thoroughly investigated in terms of size and power; simulation results exist only for a small subset of the parameter space for Dirichlet distributed data with low dimension *d*.[24] This represents are large and necessary area of potential future exploration.

While Dirichlet distribution has not been properly explored for the purpose of microbiome analysis, the Dirichlet-Multinomial distribution has been widely used for this purpose.[7, 41, 42, 43, 44, 45] ALDEx2 is a well known algorithm for microbiome analysis that utilizes a Bayesian multinomial model similar to the one shown in this work. The pure Dirichlet estimation we present here takes a fully compositional approach, that is, only information about the proportions of taxa present are retained by the model. Although modeling with the Dirichlet-multinomial distribution is considered a compositional approach, analysis is still based off of the raw counts as inputs. In this work we showed that the Dirichlet MLE achieves comparable estimates to the BDME, demonstrating the utility and potential of the Dirichlet distribution for modeling microbiomes. The framework established in this work provides a foundation for the formulation of analyses for microbiome data that are fully compositional in treatment and far more mathematically intuitive.

## 6 Acknowledgements

Daniel T. Fuller acknowledges the support from the Lawrence ‘57 and Antoinette Delaney Ignite Research Fellowship sponsored by Clarkson University, Potsdam, NY, USA.

## 7. Author contributions

DTF: Data curation, writing - initial draft, methodology, formal analysis, visualization, investigation, coding.

SM: Editing, conceptualization, methodology, investigation, project administration.

SS: Supervision, conceptualization, methodology, validation, editing.

NP: Conceptualization, methodology, validation, editing.

All authors contributed to the article and approved the submitted version.

## 8 Appendices

### 8.1 Additional simulation results

### 8.2 Additional Figures

### 8.3 Dirichlet-Multinomial Method

Let the random vector **Y** = (*Y*_1_, *Y*_2_, …, *Y*_*d*_)′ be the vector of raw observed counts from a microbiome. Typically **Y** has a multinomial distribution with a parameter vector *π* = (*π*_1_, *π*_2_, …, *π*_*d*_)′ (see, for example, Holmes, Harris, and Quince (2011)).[23] The pmf of **Y** is given as

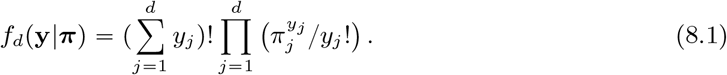

In a hierarchical Bayesian setup, let *π* ∼ Dir(***δ***|*d*) and, further, let *δ*_*j*_ ∼ Gamma(*η, ν*) with shape parameter *η* > 0 and scale parameter *ν* > 0. Let *L*(**Y**|***π***) be the multinomial likelihood, which for a single sample is equivalent to the probability mass function *f*_*d*_(**y**|***π***). The post-facto distribution of the Dirichlet parameters is

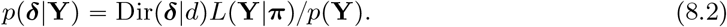

Holmes, Harris, and Quince (2012)[23] used this Bayesian model to estimate the Dirichlet parameters of the Dirichlet-Multinomial distribution for microbiome samples. While their model is intended for estimating the parameters for a mixture of Dirichlet distributions, we are interested only in the single regime case in this study. They obtained posterior parameter estimates by maximizing the evidence, which is equivalent to obtaining the posterior mode for the Dirichlet parameter vector. The evidence shown in their work simplifies to Equation 8.3, as written below.

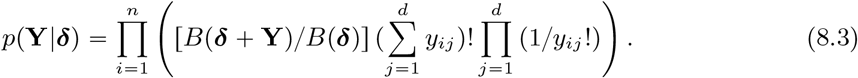

One of the limitations of this approach is that there are cases where the relative abundances are the same but the raw counts can differ. Under a Multinomial model, this can cause differences in inference. However, if an assumption is only made as relative abundances are following Dirichlet distributions regardless of any assumption on raw counts, then sample-related variations in raw abundances will be neglected.

We apply the DMM as a comparison to the Dirichlet MLE in Section 4. While the Gamma hyperparameters *η* and *ν* are not explicitly defined in the referenced work, their script sets the values at *η* = 1 and *ν* = 1 and we do the same.

## Notes

### Competing Interest Statement

The authors have declared no competing interest.

